# Validation of Duchenne muscular dystrophy candidate modifiers using a CRISPR-Cas9-based approach in zebrafish

**DOI:** 10.1101/2025.05.20.655139

**Authors:** Geremy T Lerma, Kamorah R Ryhlick, Joseph C Beljan, Olivia M Carraher, Sharon L Amacher

## Abstract

Duchenne muscular dystrophy (DMD) is a progressive muscle wasting disease for which there is no cure. There is a critical need for additional therapeutics. Human genome-wide association studies (GWAS) have identified candidate DMD genetic modifiers that could serve as therapeutic targets. Because many GWAS-identified single nucleotide polymorphisms (SNPs) lie in noncoding, putative regulatory regions, it can be challenging to identify which gene(s) are regulated by these SNPs and how gene expression is altered to modify disease severity even with extensive *in silico* modeling. We analyzed expression of zebrafish orthologs of putative DMD modifiers and showed almost all are comparably expressed in wild-type and *dmd* mutant zebrafish at three different stages of disease. To model decreased expression of candidate modifiers, we pursued a zebrafish CRISPR-based screening approach, which we validated by testing zebrafish orthologs of two extensively studied DMD modifiers, *LTBP4* and *THBS1*. We then tested candidates from the most recent GWAS and demonstrate that *galntl6, man1a1, etaa1a;etaa1b,* and *adamts17* are *bona fide* DMD modifiers. Our findings demonstrate the utility of zebrafish for DMD genetic modifier screening and characterizing modifier function.

**Summary Statement:** Zebrafish CRISPR-based screening approach validates new genetic modifiers of Duchenne muscular dystrophy.

## Introduction

Duchenne muscular dystrophy (DMD) is the most common childhood muscular dystrophy in humans and affects approximately 1 in 5,000 live male births (Emery, AE., 1991; Mendell et al., 2012; Crisafulli et al., 2020). DMD is a X-linked progressive muscle wasting disease caused by mutation of the *Dystrophin* gene that leads to complete absence of the dystrophin protein (Bushby et al., 2010). Dystrophin loss destabilizes the muscle cell membrane and leads to muscle contraction-induced injury, persistent inflammation, fibrosis, immune system dysregulation, and muscle atrophy (Bez Batti Angulski et al., 2023; Duan et al., 2021; Tidball et al., 2018). Despite development of promising DMD therapeutics (Boehler et al., 2023; D’Ambrosio & Mendell, 2023; Filippelli & Chang 2022; Guglieri et al., 2022a; Guglieri et al., 2022b; Matsuo, 2021; Wu et al., 2022), there is no cure for DMD. Additionally, there are significant challenges in Dystrophin restoration gene therapy, including patient dystrophin and AAV9 immunity (Bez batti Angulski et al., 2023; D’Ambrosio & Mendell, 2023; Mendell et al., 2010; Mendell et al., 2022). Thus, there is a critical need to identify new therapeutic targets.

Genetic modifiers are gene variants that modulate disease severity. Genetic modifiers of DMD have been identified (Flanigan et al., 2023; Wehling-Henricks et al., 2016; Weiss et al., 2018; Spitali et al., 2020; Pegoraro et al., 2011; Hogarth et al., 2017; Bello et al., 2016). Previous studies have shown that manipulating expression levels or associated pathways of known DMD modifier genes can modulate disease severity (Hogarth et al., 2017; Demonbreun et al., 2021; Kemaladewi et al., 2014; Juban et al., 2018; Ceco and McNally 2013; Granados et al., 2024). One pathway associated with known DMD modifier genes is the TGFβ signaling pathway (Todorovic and Rifkin, 2012; Adams and Lawler, 2011; Weiss et al., 2018; Flanigan et al., 2013, Heydemann et al, 2009). TGFβ signaling plays an important role in DMD pathology by regulating muscle growth and differentiation, immune response, and extracellular matrix (ECM) regulation (Ceco and McNally, 2013; Burks and Chon, 2011). In DMD, TGFβ signaling is increased (Ishitobi et al., 2000; Song et al., 2017). Increased TGFβ signaling stimulates fibrosis and inhibits muscle regeneration, exacerbating disease severity (Ceco and McNally, 2013; Burks and Cohn, 2011). Several studies have shown that inhibition of TGFβ signaling using TGFβ neutralizing antibodies or TGFβ receptor inhibitors improve DMD pathology (Burks and Cohn, 2011, Mázala et al, 2020); however, because TGFβ is involved in numerous pathways, the development of a TGFβ inhibitor with minimal adverse side effects is on-going (Massague and Sheppard, 2023). Side effects might be limited by targeting TGFβ signaling pathway regulatory proteins instead of the TGFβ receptor or ligand. For example, the TGFβ signaling pathway regulator Latent Transforming Growth Factor Protein 4 (LTBP4) was first identified as a DMD modifier via murine QTL mapping in γ-sarcoglycan–null mice (Heydemann et al, 2009), and then by candidate gene association and GWAS in human (Flanigan et al., 2013; Weiss et al., 2018). Administering anti-LTBP4 antibody improves DMD pathology and decreases TGFβ signaling in *mdx* mice and is being developed by Ikaika therapeutics for treatment of DMD and other fibrotic diseases (Demonbreun et al., 2021).

A recent GWAS study, that included the largest patient cohort to date, identified new SNPs that correlate with loss of ambulation (LoA) (Flanigan et al., 2023), all of which lie in non-coding sequence, suggesting that may be regulatory in nature (Flanigan et al., 2023). Despite extensive *in silico* modeling, it is still uncertain which gene is associated with the SNP and the direction in which expression is modulated to impact disease severity (Flanigan et al., 2023). However, regulatory variants are compelling candidates, because they suggest that gene expression modulation, instead of coding sequence alteration, is what influences DMD severity. Validation of candidate genes linked to the identified SNPs is required to confirm *bona fide* DMD modifiers.

Subsequent characterization of these *bona fide* DMD modifiers allows for insight into their mechanism of action, informs DMD pathology, and could lead to future therapeutic development.

Zebrafish, *Danio rerio*, are an established model organism to study muscular dystrophies, including DMD, and have been used in chemical and genetic DMD screens (Farr et al., 2020; Vieira et al., 2015; Johnson et al., 2013; Kawahara et al., 2011; Kawahara & Kunkel, 2013; Widrick et al., 2019). More recently, zebrafish ’crispant’ strategies have been successfully employed for genetic screening in the founder (F0) generation (Debaenst et al., 2024, Parvez et al., 2023, Trubiroha et al., 2018, Kroll et al., 2021; Shaw & Mokalled, 2021; Bek et al., 2021; Wu et al., 2018). CRISPR/Cas9-mediated mutagenesis is highly efficient and can be multiplexed to knockout gene(s) of interest (GOI); F0 crispants faithfully recapitulate genetic mutants (Kroll et al., 2021; Shaw & Mokalled, 2021; Bek et al., 2021). We utilized the advantages of the zebrafish system, including high fecundity, rapid external development, and optically transparent embryos, to perform zebrafish crispant analyses to screen putative DMD genetic modifiers identified via human GWAS. We hypothesized that if GWAS-identified SNPs decreased expression of their associated gene(s) we could mimic this effect using CRISPR-mediated knockdown. We developed a CRISPR-based screening pipeline that utilizes the birefringence properties of zebrafish muscle (Berger et al. 2012) to assess DMD onset and overall birefringence intensity. To validate our approach, we first showed that mutation of zebrafish orthologs of previously identified DMD modifiers, *LTBP4* and *THBS1* (Weiss et al., 2018), impacted DMD severity. We then tested zebrafish orthologs of 5 additional lead DMD modifier candidates, and show that mutation of 4 of them, *man1a1, galntl6, etaa1a;etaa1b,* and *adamts17*, impact disease onset, severity, and/or survival. Overall, we describe a validation method to investigate putative DMD modifiers identified via GWAS and show that *man1a1, galntl6, etaa1a;etaa1b,* and *adamts17* are *bona fide* DMD modifiers.

## Results

### Expression of candidate DMD modifiers is largely comparable in zebrafish *dmd* mutants and wild-type siblings

The 2023 GWAS conducted by Flanigan et al., identified SNPs associated with LoA in DMD that lie in non-coding, putative regulatory regions. The lead candidates for SNP-associated modifier genes were identified via extensive *in silico* modeling (Flanigan et al, 2023); however, validation is needed to confirm candidate modifier gene identity and whether the SNPs function by impacting expression of the associated modifier gene(s). We postulated that querying candidate gene expression changes across DMD progression may provide insight into how modifier SNPs alter gene expression and thus disease severity. For example, if candidate modifier transcript levels are increased in *dmd* mutants, it would suggest that modifier knockdown may impact disease severity; if transcript levels are decreased, it would suggest the opposite. We performed qPCR to quantify transcript levels of zebrafish orthologs of several lead candidates and two previously identified DMD modifiers, *LTBP4* and *THBS1*, in wild-type and *dmd*-/-mutant zebrafish at early-(4 dpf), mid-(7 dpf), and late-(10 dpf) stage disease. For some human genes, there are two zebrafish orthologs, the a and b ohnologs, that arose from the teleost whole genome duplication (Postlethwait et al., 1998). We determined the fold change between *dmd-/-* mutants and wild-type siblings (normalized to *eef1a1a*) of *thbs1a, thbs1b, ltbp4, man1a1, galntl6, etaa1a, etaa1b,* and *adamts17* at each time point (Figure 1). Only THBS1 orthologs show differences in *dmd* mutants relative to wild-type siblings, with *thbs1b* significantly increased at 4 dpf and *thbs1a* significantly increased at 7-and 10-dpf. All other tested modifiers are similarly expressed in *dmd-/-* mutants and wild-type siblings. These zebrafish expression data largely recapitulate expression trends observed for the same orthologs in 1 month old *mdx* mice (Flanigan et al., 2023). Because expression of most candidate modifiers is comparable in wild-type and *dmd* mutant zebrafish, and thus does not strongly motivate a knockdown versus over-expression screening approach, we decided to pursue the former.

**Figure 1:**
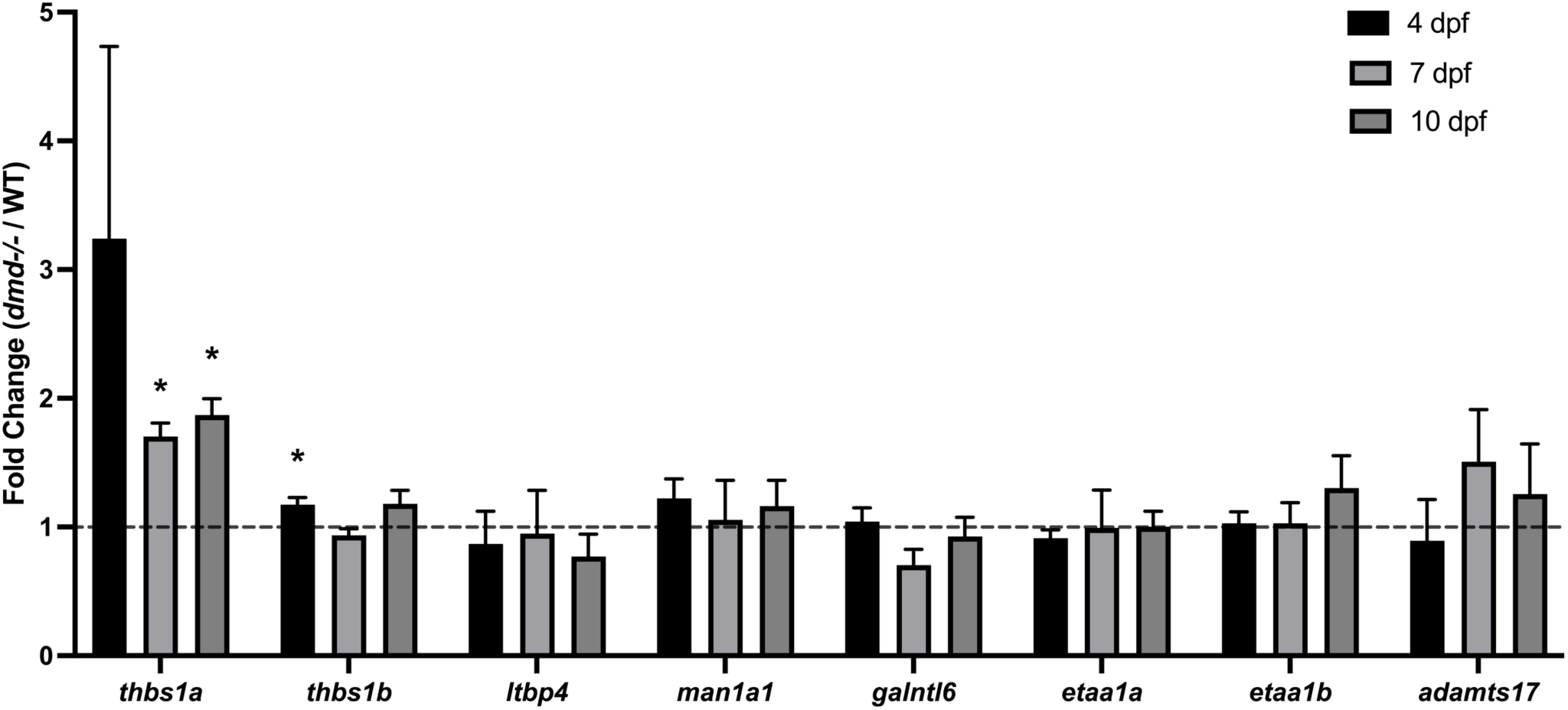
Expression of candidate DMD modifiers is largely comparable in zebrafish *dmd* mutants and wild-type siblings. Wild-type (WT) and *dmd-/-* mutants were raised to 4-(black), 7-(grey), and 10-dpf (dark grey) and processed for qPCR analysis, using 3 biological replicates (n=15 fish) for each sample and time point. qPCR data are presented as mRNA fold change in *dmd-/-* mutants relative to WT siblings of the same age, with the black dashed line indicating equivalent WT expression. *eef1a1a* was used as a normalization gene. Welch’s T-Test used, * denotes significant difference.

### Development of a Zebrafish CRISPR/Cas9 pipeline to assess functional impact of DMD modifier candidates

We developed a zebrafish CRISPR-based strategy to assess whether knockdown of candidate modifier expression influences DMD phenotypes (Figure 2A). For each candidate modifier, embryos from a *dmd+/-* intercross are microinjected with Cas9 protein and CRISPR(s) targeting the candidate gene or raised as uninjected controls. For candidate modifier genes with two zebrafish orthologs, we targeted both copies of the duplicated gene to avoid genetic compensation (El-Brolosy & Stainier, 2017). Because DMD in zebrafish follows an autosomal recessive inheritance pattern, approximately 25% of the intercross progeny are *dmd-/-* and the remaining 75% are phenotypically wild-type (Bassett et al., 2003), allowing us to compare the effect of modifier knockdown in *dmd* mutants and wild-type siblings, using birefringence to determine whether modifier loss of function affects DMD onset and/or overall muscle organization (Figure 2A). Birefringence imaging is a non-invasive imaging technique that relies on the highly ordered crystalline-like structure of zebrafish muscle (Berger et al., 2012). When two polarizing filters are aligned perpendicular to one another, very little light is transmitted. However, placing a highly ordered crystal-like sample between the polarizers causes light diffraction called birefringence (Berger et al., 2012). The highly ordered muscle structure in wild-type zebrafish cause bright birefringence, whereas disorganized muscle, like dystrophic lesions present in *dmd-/-* mutant fish, are non-birefringent and appear as dark patches (Figure 2B,B’) (Berger et al., 2012). The DMD onset assay is a qualitative assay that determines the day on which a live fish first displays any non-birefringent muscle integrity anomaly, regardless of severity (Figure 2B,B’). To quantitatively assess muscle integrity, we analyzed total birefringence in fixed specimens at 4 dpf, the day on which all homozygous *dmd* mutants from a *dmd+/-* intercross display muscle integrity defects. We note that day of onset does not correlate with birefringence intensity at 4 dpf (R^2^ = .02).

**Figure 2:**
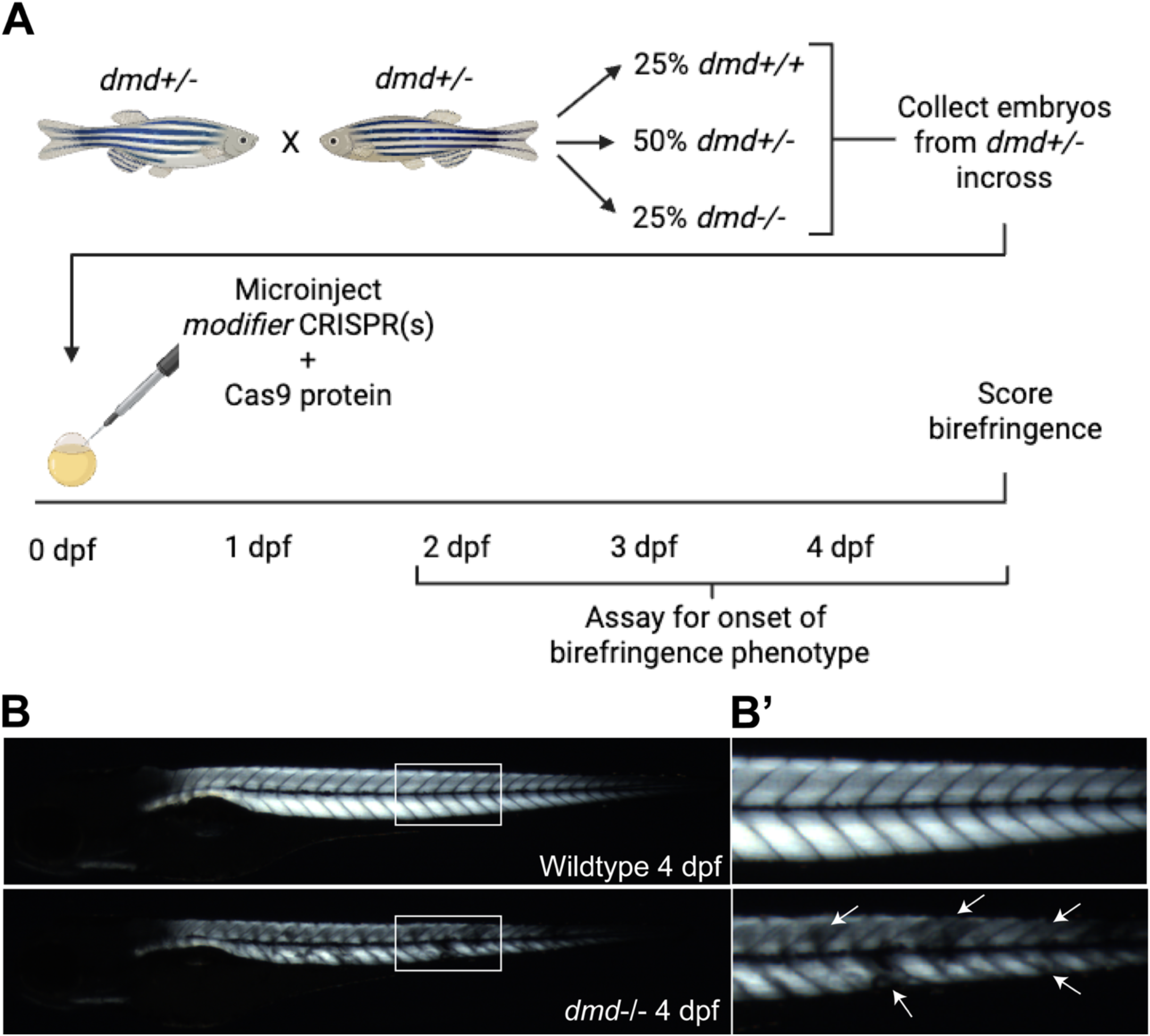
**Development of a Zebrafish CRISPR/Cas9 pipeline to assess functional impact of DMD modifier candidates**. (A) Adult fish heterozygous for *dmd* are intercrossed to generate embryos for the screen. Because DMD in zebrafish follows an autosomal recessive inheritance pattern, one-quarter of the intercross progeny are homozygous *dmd* mutants (*dmd-/-*), all of which are identifiable using birefringence imaging by 4 dpf due to the presence of muscle lesions. Homozygous wild-type (*dmd+/+*) and heterozygous *dmd* (*dmd+/-*) fish are indistinguishable. Embryos are injected by the 1-cell stage with modifier-targeting CRISPR and Cas9 protein or raised as uninjected controls. All fish are assayed for DMD onset between 2-4 dpf using live birefringence imaging. At 4 dpf, all fish are fixed, birefringence imaged, and genotyped via HRMA. (B, B’) Representative examples of wild-type (WT, top) and *dmd-/-* mutant (bottom) fish imaged via birefringence at 4 dpf. The boxes in B are shown magnified in B’. Wild-type fish show strong, uniform birefringence, whereas *dmd* mutant fish have variably-sized non-birefringence patches indicating dystrophic lesions (white arrows). Fish containing one or more lesion, regardless of lesion size or number, are scored as dystrophic in the onset assay.

Because human SNPs likely modulate gene expression of the candidate gene, we hypothesized that complete loss of function would not be required to observe an effect. To assess mutagenesis efficiency, we performed high-resolution melt analysis (HRMA) of DNA from every crispant phenotyped in the pipeline. While HRMA does not measure indel frequency directly, it has been used to reliably estimate successful mutagenesis (Talbot & Amacher, 2014; Dahlem et al., 2012; Samarut et al., 2016). The data show that CRISPR/Cas9 microinjections targeting 2 known modifiers (3 genes) and 6 candidate modifiers (8 genes) led to an average of 94% mutagenesis detection via HRMA across all experiments (Fig. S1A, B).

### Mutation of previously identified DMD modifiers *LTBP4* and *THBS1* impacts DMD severity in zebrafish

A previous human GWAS identified variant alleles of two TGFβ regulator genes, *LTBP4,* that encodes Latent TGFβ Binding Protein 4, a protein that sequesters TGFβ latency-associated peptide complex (LAP), and *THBS1*, that encodes Thrombospondin 1, a multifunctional ECM glycoprotein that activates TGFβ by releasing it form the LAP, as DMD modifiers (Weiss et al., 2018). The protective modifier alleles caused decreased LTBP4 and THSB1 expression (Weiss et al., 2018), respectively, thus these genes were ideal proof-of-principle loci to validate our screening approach. Before generating and analyzing crispants, we first assessed whether inhibiting TGFß signaling, the pathway in which both LTBP4 and THBS1 function, modulates the DMD phenotype in zebrafish, as it does in other animal models (Ceco and McNally, 2013; Kemaladewi et al, 2014). We treated progeny of a *dmd+/-* intercross with a TGFβ inhibitor, SB431542 (Inman et al., 2002; Callahan et al., 2002) Because prolonged TGFβ inhibition leads to developmental defects (data not shown), we used a short 24-hour treatment window between 24-48 hpf that encompasses the time during which muscle lesions first appear. We found that TGFβ inhibition with SB431542 significantly increases birefringence intensity (Fig. S2), further motivating testing of LTBP4 and THBS1 orthologs in our screening pipeline.

We targeted zebrafish orthologs of *LTBP4 (ltbp4)*, and *THBS1* (*thbs1a* and *thbs1b)* (Figure 3). DMD onset is unaffected in *dmd-/-;ltbp4* crispants (Figure 3A), and birefringence intensity is significantly increased (Figure 3B, C), suggesting improvement in muscle organization. Birefringence intensity is also significantly improved in *dmd-/-;thbs1a;thbs1b* crispants, although DMD onset is accelerated (Figure 3D-F). Because the longest ambulating patient from the Weiss et al (2018) study was homozygous for both *LTBP4* and *THBS1* protective alleles, we tested the combinatorial impact of targeting both zebrafish orthologs. Like the *dmd-/-;thbs1a;thbs1b* crispants, the *dmd-/-;ltbp4;thbs1a;thbs1b* crispants display significantly increased birefringence intensity and accelerated DMD onset (Figure 3 G-I). Although it was surprising that DMD onset was accelerated in the latter two cases, we note that the onset assay only assesses the first appearance of a muscle lesion, not phenotypic severity. The improvement of overall birefringence, which does assess severity, supports that our screening approach can identify genetic modifiers of DMD.

**Figure 3:**
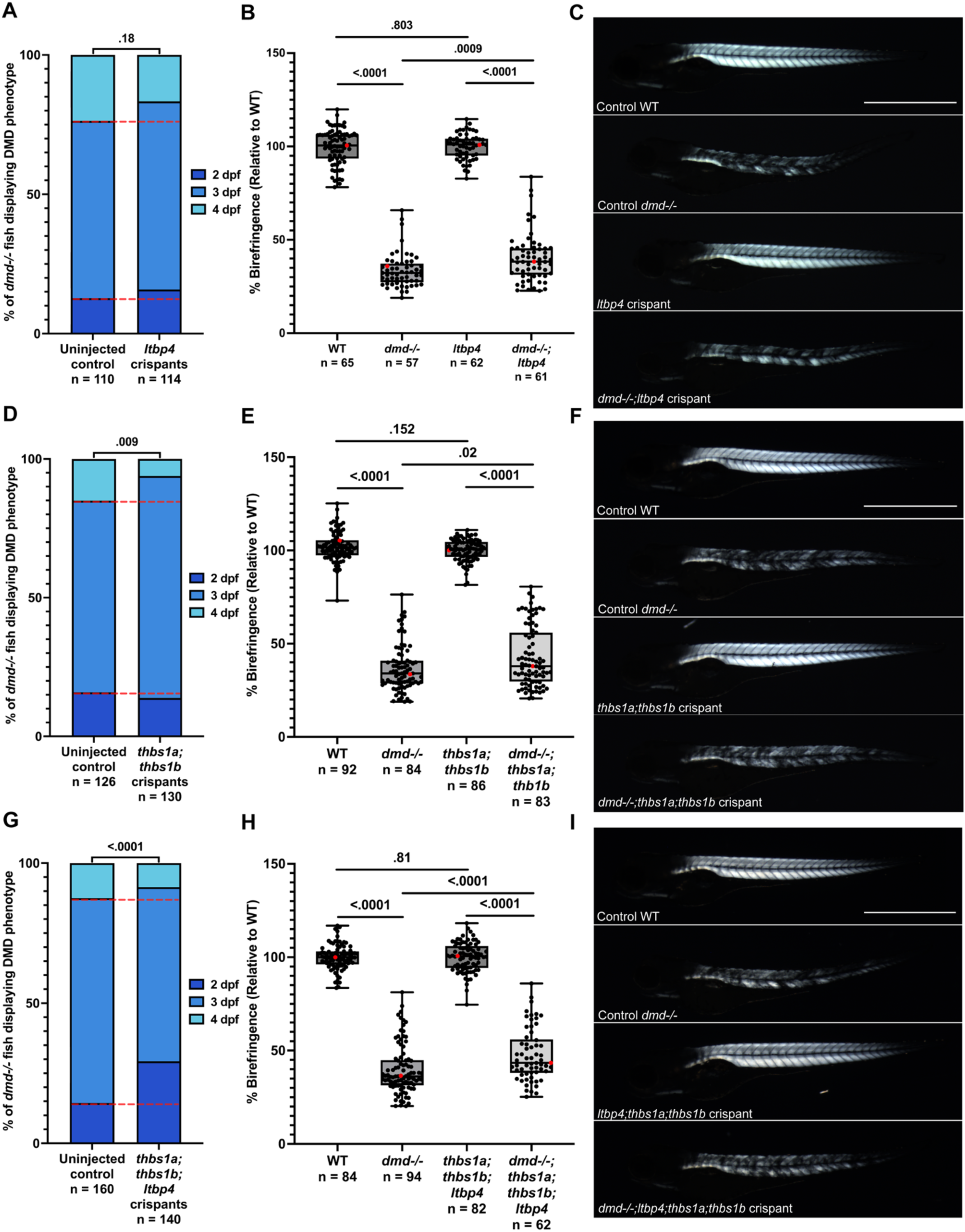
Mutation of previously identified DMD modifiers *LTBP4* and *THBS1* impacts DMD severity in zebrafish. Embryos from a *dmd+/-* intercross were injected with *ltbp4-* (A-C), *thbs1a- and thbs1b-* (D-F), or *ltbp4-, thbs1a-,* and *thbs1b-* (G-I) targeting CRISPR(s) and Cas9 protein or raised as uninjected controls. (A,D,H) DMD onset was assessed at 2 dpf (dark blue), 3 dpf (blue), and 4 dpf (light blue). Chi-square analysis was carried out to determine if there was a significant different in onset age proportions. (B,E,I) Box plots show overall birefringence intensities at 4 dpf. Mann-Whitney U-Tests carried out for comparisons of overall birefringence intensities. Red dots denote representative images shown in panels C, F, J. (C,F,J) Representative birefringence images for each condition. Scale bar denotes 1mm. Experiments were conducted at minimum in biological triplicate. Sample sizes listed below each condition.

### Mutation of *galntl6* accelerates DMD onset and mutation of *man1a1* improves birefringence intensity

Two of the lead candidate modifier genes, *GALNTL6* and *MAN1A1*, encode enzymes required for various steps in glycosylation. The top candidate modifier gene for GWAS-identified SNP rs1358596 is *GALNTL6* (Flanigan et al, 2023), that encodes polypeptide N-acteylgalactosaminyltransferase like 6, a protein that catalyzes the initial reaction in O-linked glycosylation (Raman et al, 2012; Peng et al, 2010). Another GWAS correlated *GALNTL6* with strength versus endurance disposition in athletes (Diaz Ramirez et al., 2020). We targeted *galntl6*, the zebrafish *GALNTL6* ortholog, and observed that DMD onset is significantly accelerated (Figure 4A) whereas birefringence intensity is comparable (Figure 4B, C) in *dmd-/-;galntl6* crispants compared to *dmd-/-* siblings. The top candidate modifier gene for GWAS-identified SNP rs10499096 is *MAN1A1* (Flanigan et al, 2023), that encodes mannosidase alpha class 1A member 1, a protein involved in removal of α1,2 linked mannose structures (Jin et al., 2018). We targeted *man1a1*, the zebrafish *MAN1A1* ortholog, and observed that DMD onset is comparable (Figure 4D) and birefringence intensity is significantly improved (Figure 4E,F) in *dmd-/-;man1a1* crispants compared to *dmd-/-* siblings. These data support that *galntl6* and *man1a1* are *bona fide* DMD modifiers.

**Figure 4:**
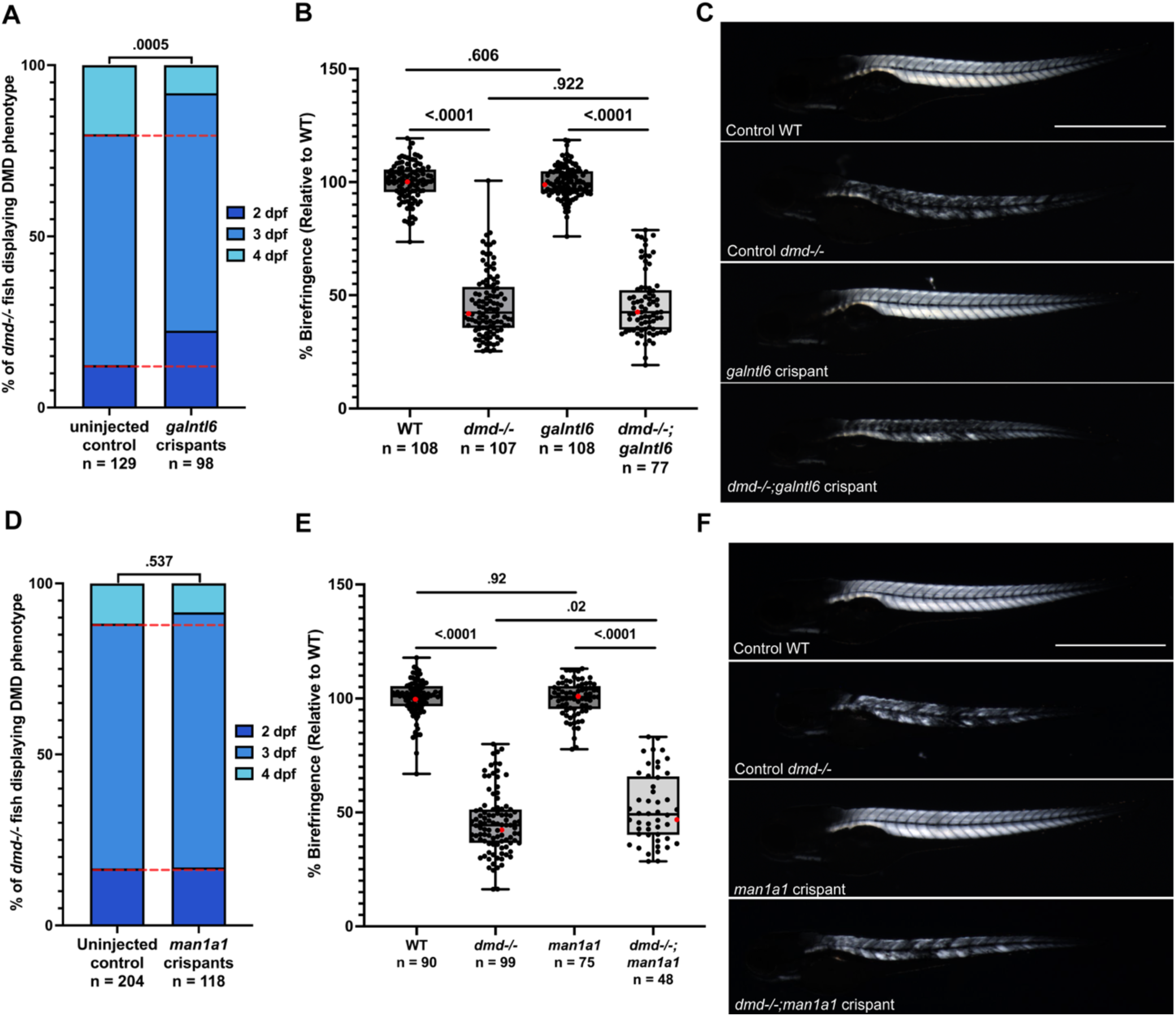
Mutation of *galntl6* accelerates DMD onset and mutation of *man1a1* improves birefringence intensity. Embryos from a *dmd+/-* intercross were injected with *galntl6* (A-C) or *man1a1* (D-F) targeting CRISPR and Cas9 protein or raised as uninjected controls. (A,D) DMD onset was assessed at 2 dpf (dark blue), 3 dpf (blue), and 4 dpf (light blue). Chi-square analysis was carried out to determine if there was a significant difference in onset age proportions. (B,E) Box plots show overall birefringence intensities. Mann-Whitney U-Tests carried out for comparisons of overall birefringence intensities. Red dots denote representative images shown in panels C and F. (C,F) Representative birefringence images for each condition. Scale bar denotes 1mm. Experiments were conducted at minimum in biological triplicate. Sample sizes listed below each condition.

### Mutation of *etaa1a* and *etaa1b* delays DMD onset

The top candidate modifier genes for GWAS-identified SNPs rs34263553 and rs2061566, *ETAA1* and *PARD6G*, encode proteins involved in various aspects of stem cell proliferation and regulation (Bass et al, 2016; Misoge et al, 2017; Bass & Cortez, 2019; Feige et al, 2018; Yamashita et al, 2010). ETAA1 encodes ETAA1 activator of ATR (Serine/threonine-protein) kinase, a protein regulating replication stress and DNA damage response pathways (Bass et al., 2016). PARD6G encodes Par-6 family cell polarity regulator gamma, a protein involved in cell polarity maintenance (Dumont et al., 2015). Because mutation of zebrafish *pard6gb* causes a strong developmental phenotype (Grant & Moens, 2010), we did not screen the zebrafish orthologs *pard6ga* and *pard6gb* in our crispant pipeline. We did pursue *ETAA1* orthologs, *etaa1a* and *etaa1b*, and found that DMD onset is significantly delayed (Figure 5A) and birefringence intensity is increased (almost significantly, p=0.051) (Figure 5B,C) in *dmd-/-;etaa1a;etaa1b* crispants compared to *dmd-/-* siblings. These data support that *etaa1a* and *etaa1b* are *bona fide* DMD modifiers.

**Figure 5:**
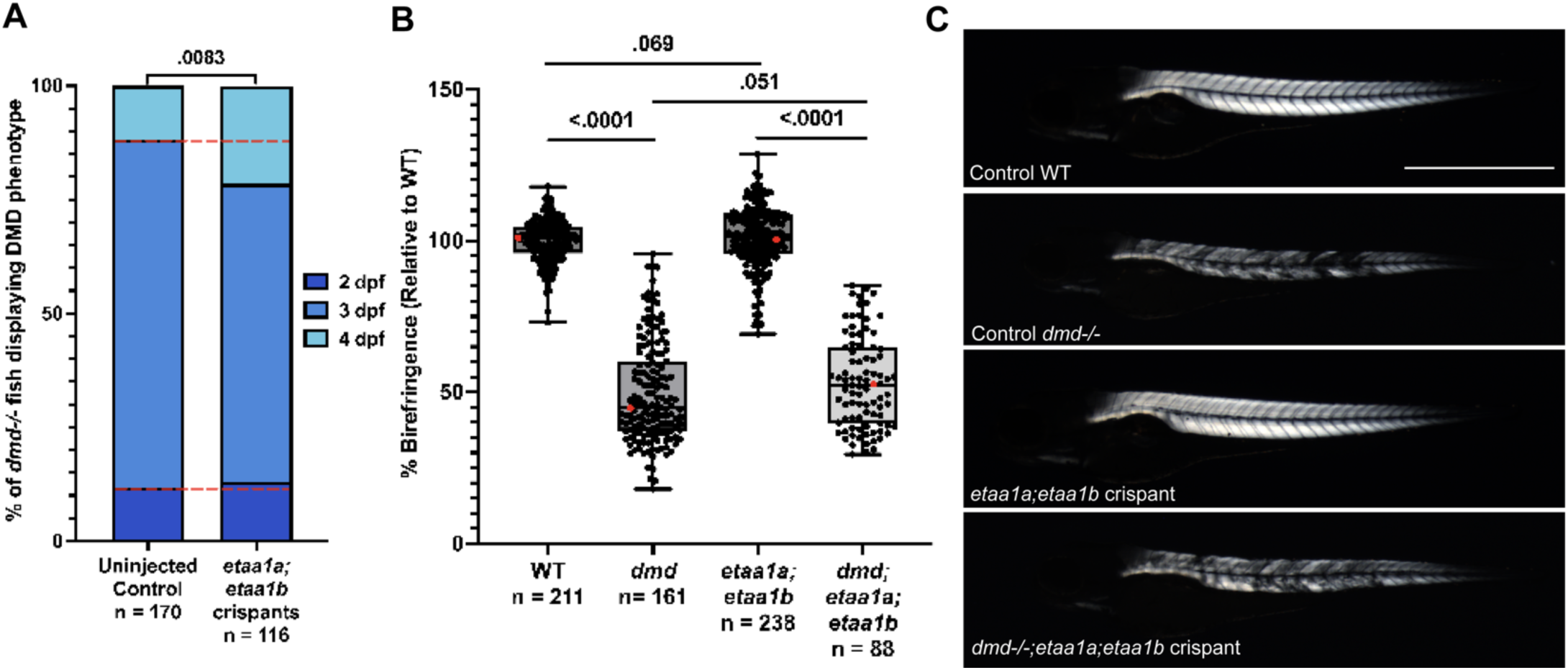
Mutation of *etaa1a* and *etaa1b* delays DMD onset. Embryos from a *dmd+/-* intercross were injected with *etaa1a* and *etaa1b* targeting CRISPRs and Cas9 protein or raised as uninjected controls. (A) DMD onset was assessed at 2 dpf (dark blue), 3 dpf (blue), and 4 dpf (light blue). Chi-square analysis was carried out to determine if there was a significance different in onset age proportions. (B) Box plots show overall birefringence intensities. Mann-Whitney U-Tests carried out for comparisons of overall birefringence intensities. Red dots denote representative images shown in panel C. (C) Representative birefringence images for each condition. Scale bar denotes 1mm. Experiments were conducted at minimum in biological triplicate. Sample sizes listed below each condition.

### Mutation of *adamts17* delays DMD onset, improves birefringence intensity, and improves survivorship of *dmd* mutants

The top candidate modifier gene for GWAS-identified SNP rs10077875 is *ADAMTS19*, which encodes ADAM metallopeptidase with thrombospondin type 1 motif 19, a protein involved in ECM remodeling. This gene has been lost in most teleosts, including zebrafish, however zebrafish do have an ortholog of *ADAMTS17*, a closely related sister gene of *ADAMTS19* that arose from a vertebrate genome duplication event (Kelwick et al., 2015). Protein alignment shows 55.7% overall identity between human ADAMTS17 and ADAMTS19, 55.8% between human ADAMTS19 and zebrafish Adamts17, and 69.6% between human ADAMTS 17 and zebrafish Adamts17 (Fig. S3A). ADAMTS proteins contain two major domains: a catalytic domain (peptidase M12B and disintegrin) and an ancillary domain (five TSP1 repeats, spacer, and PLAC). Between human ADAMTS19 and zebrafish Adamts17, the catalytic domains share 65.9% (peptidase M12B) and 64.6% (disintegrin) identity (82.3% and 83.1% similarity), while the ancillary TSP1 domains share 49.72% identity (63.04% similarity) (Fig. S3B, C) and manual comparisons of the ancillary spacer and PLAC domains shows 56.29% and 87.5% identity (67% and 97% similarity), respectively.

*ADAMTS17* and *ADAMTS19* are broadly expressed in humans (Uhlen M, et al, 2015; Karlsson M, et al, 2021), with overlapping and distinct sites of expression that correlate with distinct phenotypes observed in patients with the corresponding human autosomal recessive disorders. *ADAMTS19* is more highly expressed than *ADAMTS17* in the heart, and autosomal recessive mutation of *ADAMTS19* leads to Cardiac valvular dysplasia 2 (Wunnemann et al, 2020; Massadeh et al, 2020; Uhlen M, et al, 2015; Karlsson M, et al, 2021). *ADAMTS17* is more broadly expressed than *ADAMTS19*, including in the pituitary gland and the skin where *ADAMTS19* is not expressed, and is more highly expressed in the retina than *ADAMTS19* (Uhlen M, et al, 2015; Karlsson M, et al, 2021). Homozygous mutation of *ADAMTS17* leads to Weill-Marchesani syndrome 4, an autosomal recessive connective tissue disorder characterized by short stature, brachydactyly, small spherical and dislocated lenses, thick skin, and occasionally a heart valve defect (Evans et al, 2020; Shah et al, 2014). The specific phenotypes caused by *ADAMTS17* and *ADAMTS19* loss of function suggest that the two genes compensate for each other in tissues where they are co-expressed. Zebrafish *adamts17* is expressed in all tissues where *ADAMTS19* and *ADAMTS17* are expressed in humans, with highest expression in skeletal muscle, heart, and gut (Brunet et al, 2015). These observations, together with previous studies that have suggested ADAMTS17 and ADAMTS19 work together in a network to regulate ECM dynamics (Karoulias et al, 2020), suggest that the single zebrafish *adamts17* gene performs the functions that both *ADAMTS17* and *ADAMTS19* perform in humans.

We thus targeted *adamts17* and found that DMD onset is delayed (Figure 6A) and birefringence intensity is increased (Figure 6B,C) in *dmd-/-;adamts17* crispants compared to *dmd-/-* siblings. Because both parameters indicate improvement upon *adamts17* mutation, we also assessed survivorship. We raised wild-type, *dmd-/-*, *adamts17* crispant, and *dmd-/-;adamts17* crispant fish and tracked survivorship through 12 dpf (Figure 6D). Wild-type and *adamts17* crispant fish have similarly high survivorship. Both *dmd-/-* mutants and *dmd-/-;adamts17* crispants have significantly lower survivorship than wild-type and *adamts17* crispant fish; however, survivorship is significantly improved in *dmd-/-;adamts17* crispants compared to *dmd-/-* siblings (Figure 6D). Together, these data strongly support that *adamts17* is a *bona fide* DMD modifier.

**Figure 6:**
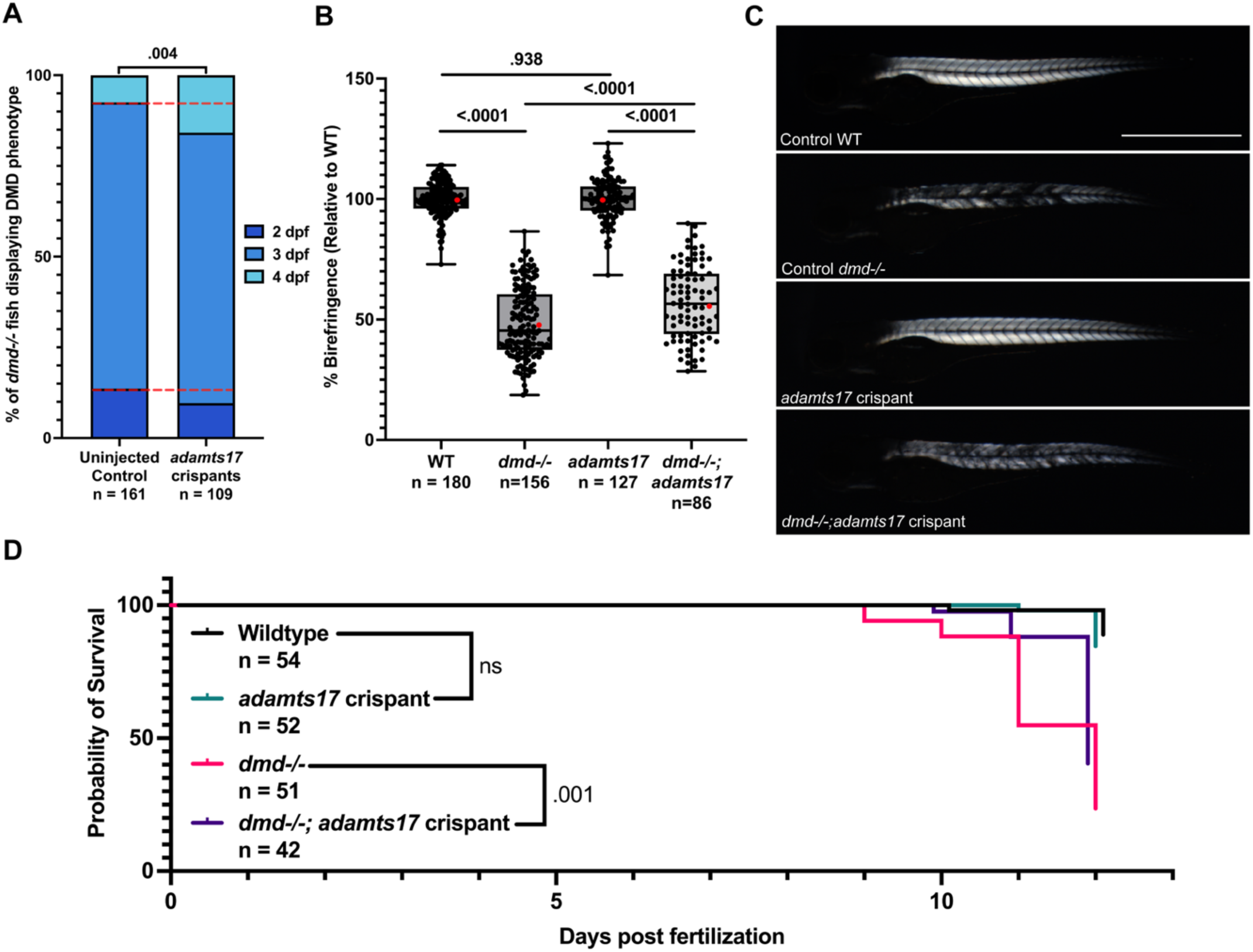
Mutation of *adamts17* delays DMD onset, improves birefringence intensity, and improves survivorship of *dmd* mutants. Zebrafish do not have a homolog of *ADAMTS19*. To test the effect of *adamts17* mutation on DMD embryos from a *dmd+/-* intercross were injected with *adamts17* targeting CRISPR and Cas9 protein or raised as uninjected controls. (A) DMD onset was assessed at 2 dpf (dark blue), 3 dpf (blue), and 4 dpf (light blue). Chi-square analysis was carried out to determine if there was a significant difference in onset age proportions. (B) Box plots show overall birefringence intensities. Mann-Whitney U-Tests carried out for comparisons of overall birefringence intensities. Red dots denote representative images shown in panel C. (C) Representative birefringence images for each condition. (D) Standard survivorship assays were performed between 0-12 dpf with wild-type (black), *adamts17* crispants (green), *dmd-/-* (pink), and *dmd-/-;adamts17* crispants (purple). Log-rank test for trend was carried out: wild-type vs *dmd-/-* pvalue = <.0001 and *adamts17* crispant vs *dmd-/-;adamts17* crispant pvalue = <.0001. Experiments conducted at minimum in biological triplicate. Sample sizes listed below each condition.

### Mutation of zebrafish *NCALD* and *KLF10* orthologs does not affect DMD phenotype

We have shown that decreased function of four of six top candidate DMD modifier genes identified by Flanigan et al (2023) impact DMD onset, severity, or both. As mentioned previously, we did not test *PARD6G* orthologs due to the previously characterized zebrafish *pard6gb* mutant phenotype that would confound birefringence phenotyping. The sixth top candidate modifier gene, associated with SNP rs72681143, is *NCALD*, that encodes Neurocalcin delta, a protein involved in calcium sensing that has been shown to regulate cyclic GMP synthesis (Duda et al, 2018) and has been previously identified as a modifier of spinal muscular atrophy (Riessland et al, 2017).

Transcripts of the two zebrafish *NCALD* orthologs, *ncalda* and *ncaldb*, are expressed similarly in *dmd-/-* mutants and wild-type siblings at 4, 7, and 10 dpf via qPCR, with a slight, but significant, decrease in *ncaldb* expression at 4 dpf (Fig. S4A). We targeted *ncalda* and *ncaldb* in our crispant screen and observed no significant difference in either DMD onset or birefringence intensity in *dmd-/-;ncalda;ncaldb* crispants compared to *dmd-/-* siblings (Fig. S4B-D). Although Flanigan et al (2023) selected *NCALD* as the lead candidate for the rs72681143SNP, they suggested that another linked gene, *KLF10*, was also a credible candidate. *KLF10* encodes the protein Kruppel-like factor 10 that is known to modulate TGFß signaling (Subramaniam et al, 1995; Spittau and Krieglstein, 2012) and has been previously shown to modify DMD in *mdx* mice (DiMario, 2018). Thus, we also tested *klf10* as a possible DMD modifier. Expression of the zebrafish ortholog, *klf10*, is generally higher in *dmd-/-* mutants compared to wild-type siblings at all stages examined, however differences are not statistically significant (Fig. S4A). We targeted *klf10*, and observed no significant difference in either DMD onset or birefringence intensity in *dmd-/-;klf10* crispants compared to *dmd-/-* siblings (Fig. S4E-G). These data support that neither *ncalda/ncaldb* nor *klf10* are DMD modifiers, although it is possible that they may influence disease in combination, with other modifiers, or with standard of care treatment.

### A CRISPR-based screening approach in zebrafish validates candidate human GWAS-identified DMD modifiers

In Figure 7, we summarize the effect of knockdown of each candidate modifier(s) on the major DMD phenotypes we screened in this study. To assess global differences in birefringence intensity among *dmd-/-;*crispant groups tested, we calculated the percent difference in birefringence intensity (Figure 7A). To compare the shifts in disease onset, we compared the percent difference in the day of onset proportions for each *dmd* mutant GWAS modifier combination to their respective non-crispant control (Figure 7B). These comparisons reveal that mutation of 4 human gene orthologs (*ltpb4*, *thbs1a/thbs1b*, *man1a1*, and *adamts17*) all improve birefringence intensity by about 25% and mutation of 4 human gene orthologs (*thbs1a/thbs1b*, *galntl6*, *etaa1a/etaa1b*, and *adamts17*) either accelerate or delay disease onset. When considered together, *adamts17* emerges as a strong modifier gene as it delays onset, improves muscle integrity, and improves survivorship of *dmd-/-* mutants.

**Figure 7:**
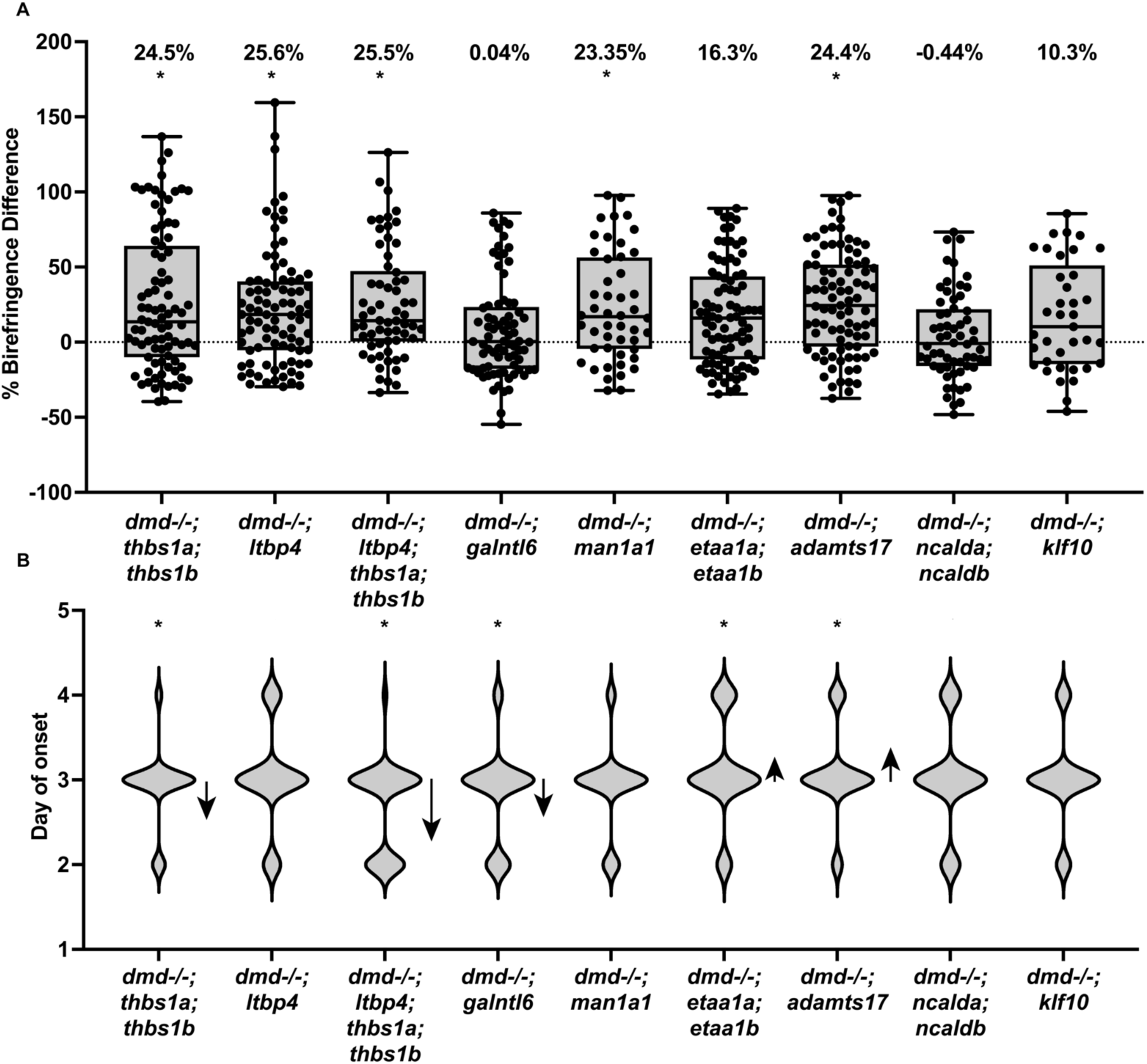
Summary of *modifier* mutation effects on DMD birefringence and onset. Analysis of the trends in shifts in overall birefringence and DMD onset for each *dmd-/-;crispant* combination. A) Percent birefringence difference was calculated by subtracting the relative birefringence intensity for each crispant modifier group to the median of their respective control and dividing by the control median intensity and expressing the result as a percentage. Average percent difference listed on top, * denotes significance via Mann-Whitney U-Tests shown in previous figures. B) Violin plots showing the proportion of fish first displaying a DMD phenotype at the specified time point for each genotype. * denotes significance via chi-square analysis shown in previous figures. Arrows represent trend change (earlier versus later onset); arrow length is proportional to the magnitude of the total change.

## Discussion

In this study, we present a zebrafish CRISPR-based screening pipeline to assess whether knockdown of putative DMD modifiers influence DMD phenotypes. We validate our approach by demonstrating that knockdown of previously identified DMD modifiers, *ltbp4* and *thbs1a* influence DMD onset and overall birefringence. We provide evidence that 4 DMD modifier candidates identified via human GWAS, *man1a1*, *galntl6*, *etaa1a;etaa1b*, and *adamts17,* are *bona fide* DMD modifiers. Our work builds upon DMD GWAS that identify putative DMD modifiers by providing a validation pipeline to identify and prioritize lead candidates for further investigation and for validation in other models.

### Expression levels of putative DMD modifiers is largely comparable between WT and *dmd* mutant zebrafish

Many human GWAS identified variant SNPs lie in non-coding, putative regulatory regions. Because such SNPs are located outside the coding region, it is typically assumed that they act by influencing the expression and/or levels of the associated modifier gene product(s). Alterations of modifier gene expression may be a normal part of disease pathology; if so, then a variant SNP may alleviate or aggravate pathology by directly modulating expression of the modifier. For example, SPP1 is a multifunctional cytokine involved in inflammation, mineralization, and immune cell modulation (Lund et al., 2009; Zhao et al., 2024; Icer and Gezmen-Karadag, 2018; Kramerova et al., 2019). SPP1 expression is elevated in human DMD patients and *mdx* mice (Zanotti et al., 2011; Capote et al., 2016; Vetrone et al., 2009). Previous studies have demonstrated that decreased Spp1 expression in *mdx* mice ameliorates DMD phenotypes (Capote et al., 2016; Vetrone et al., 2009 Kramerova et al., 2019). In humans, glucocorticoid treatment in individuals with the variant SPP1 SNP rs28357904 correlates with increased SPP1 expression, decreased grip strength, and earlier loss of ambulation (Pegoraro et al, 2011; Bello et al, 2012; Bello and Pegoraro, 2019). Although not directly comparable, these data illustrate modulation of gene expression modify disease pathology.

Alternatively, expression of a modifier gene may not normally change over the course of the disease, but a SNP that causes modifier over- or under-expression may impact disease pathology by influencing activity or expression of other components. For example, TGFß signaling is elevated in DMD patients (Ishitobi et al., 2000; Song et al., 2017), and variant SNPs that alter expression of TGFß signaling pathway modulatory genes, such as *LTBP4*, are known DMD modifiers (Weiss et al., 2018). Flanigan et al (2023) reported that, except for *THBS1*, expression of lead candidate modifiers identified by non-coding SNPs in two GWAS (Weiss et al, 2018, Flanigan et all, 2018) were not significantly different in Δ51 *mdx* mice compared to wild-type mice at 1 month, which is consistent with the expression data we present here for the orthologous genes in zebrafish (Figure 1). These data suggest that expression levels of most of the candidate modifiers we tested in our screening pipeline are not changing over the course of the disease.

### Design of a zebrafish CRISPR-based screen to identify genetic modifiers of DMD

As mentioned above, human GWAS-identified non-coding SNPs are thought to impact expression of their associated gene(s). We hypothesized that if the SNP leads to decreased expression of the associated gene(s) then CRISPR/Cas9 mutagenesis would mimic the SNP effect. Zebrafish CRISPR/Cas9 ‘crispant’ screens have been previously shown to be highly efficient in inducing mutagenesis of the targeted gene (Debaenst et al, 2024, Parvez et al, 2023, Trubiroha et al, 2018, Kroll et al, 2021; Shaw & Mokalled, 2021; Bek et al, 2021). We adapted the crispant screening approach to target zebrafish orthologs of putative DMD modifiers to test if modifier mutation impacts DMD onset or disease severity using the birefringence assay (Figure 2). HRMA analysis demonstrates that mutagenesis of targeted genes is highly efficient, on average inducing mutagenesis in 94% of injected embryos, and that multiplexing (up to 3 guide RNAs) does not impact efficiency (Fig. S1). Using our screening strategy, we validated 4 novel DMD modifier genes: *galntl6, man1a1, etaa1a;etaa1b, and adamts17*. Our efficient and high-throughput CRISPR-based screening strategy for disease modifiers should be adaptable to other human diseases that can be modeled in zebrafish.

Although we focused on knockdown to assess impact of loss of modifier function on disease pathology, over-expression approaches could also be integrated into the strategy to assess the effect of boosting modifier expression on disease pathology.

### Like in mammals, *LTBP4* and *THBS1* are also DMD modifiers in zebrafish

LTBP4 and THBS1 are regulators of TGFβ that have been identified as human DMD genetic modifiers (Todorovic and Rifkin, 2012; Adams and Lawler, 2011; Weiss et al., 2018; Flanigan et al., 2013). TGFβ levels are elevated in DMD patients (Ishitobi et al, 2000; Song et al, 2017), and previous studies have demonstrated that TGFβ signaling inhibition ameliorates DMD severity (Ceco and McNally, 2013; Kemaladewi et al, 2014). Similarly, we show that treating *dmd-/-* zebrafish with TGFβ inhibitors SB431542 improves birefringence intensity (Fig. S2). However, prolonged TGFβ inhibition is not a viable therapeutic strategy due to TGFβ’s pleiotropic roles in development (Massague, 2012; Burks and Chon, 2011; Delaney et al., 2017; Yu et al., 2017). However, modulating TGFβ signaling through TGFβ regulatory proteins may ameliorate disease severity while also minimizing adverse effects. For example, treatment with an anti-LTBP4 antibody in *mdx* mice reduces fibrosis and increases myofiber size (Demonbreun A.R, et al, 2021) and is now being developed by Ikaika Therapeutics as a therapeutic for the treatment of fibrotic diseases. Developing a rapid pipeline to validate, identify, and characterize DMD modifiers will help prioritize candidates for therapeutic development.

We selected zebrafish *LTBP4* and *THBS1* orthologs as proof-of-principle genes for our screening strategy. We show that loss of *ltbp4* function and combined loss of *thbs1a* and *thbs1b* function increases birefringence intensity (Figure 3) and thus ameliorates DMD severity. Because the longest ambulating patient from Weiss et al, 2018 was homozygous for decreased expression alleles of <u>both</u> *LTBP4* and *THBS,* we also tested whether combinatorial loss of *ltbp4*, *thbs1a*, and *thbs1b* showed an even larger improvement, and observed that *dmd-/-;ltbp4;thbs1a;thsb1b* crispants displayed the same increased birefringence intensity as in *dmd-/-;ltbp4* crispants and *dmd-/-;thbs1a;thsb1b* crispants at 4 dpf, during early stages of the disease (Figures 3, 8). Surprisingly, we observe that DMD onset is accelerated in *dmd-/-;thbs1a;ths1b* crispants and *dmd-/-;ltbp4;thbs1a;thbs1b* crispants, but not *dmd-/-;ltbp4* crispants (Figures 3, 8), suggesting that *ltbp4* is indeed a strong candidate therapeutic target. TGFß signaling plays critical roles in muscle development and in ECM regulation (Hinz, B. 2015; Ceco & McNally, 2013; Ismaeel A, *et al.,* 2019; Delaney et al., 2016). It is likely that the highly efficient knockdown achieved by our approach is greater than that caused by the human non-coding variants, and thus interferes with some normal functions of TGFß signaling in addition to pathological effects of elevated TGFß signaling.

### *man1a1* as a modifier of DMD

*MAN1A1* belongs to a family of 3 mannosidases that remove α1,2-linked mannose sugars from high mannose glycans, allowing for the formation of hybrid and complex N-linked structures (Legler et al, 2018). Disruption of MAN1A1 and MAN1A1 family members increases high mannose glycan concentration at the expense of hybrid and complex N-linked glycans (Jin et al, 2018). N-linked glycans are involved in numerous processes including of facilitation of cell-cell adhesion, immune system regulation, muscle ion function, and myogenesis (Zhang et al, 1999; Esmail and Manolson 2021; Blazev et al, 2021; Varki 2017; Pinho et al, 2023). In DMD, perturbations of the immune system, myogenesis, and ECM dynamics play key roles in disease severity modulation (Duan et al, 2021). Macrophages and dendritic cells express C-type lectin receptors which recognize mannose-, fucose-, N- acetylgalactosamine-, and N-acetylclucosamine-containing glycoconjugates, whose abundance may be shifted upon loss of MAN1A1, thus perturbing immune response (Geijtenbeek et al., 2009). In myogenesis, N-linked glycosylation is heavily regulated (Blazev et al, 2021). Gene ontology analysis of genes upregulated during rodent myogenesis shows enrichment for genes involved in N-glycan biosynthesis, specifically those involved in α-2, α-3 sialylation (Blazev et al, 2021). Loss of MAN1A1 leads to the decrease of complex N-linked structures such as sialic acids (Jin et al, 2018). KLOTHO is a protein with α2,6 sialidase activity that reduces cell surface sialic acids (Kurosu et al, 2005). Following acute muscle injury, *Klotho* undergoes muscle specific silencing (Welc et al., 2020, Wehling-Hendricks et al., 2016). It has been demonstrated that transgenic expression of *klotho* in the *mdx* mouse improves disease pathology (Wehling-Hendricks et al., 2016) and that loss of *klotho* expression impairs muscle stem cell function and muscle generation in mice (Ahrens et al., 2018). Additionally, muscle sodium channels are heavily glycosylated with a large portion of being sialic acids (Zhang et al., 1999; Bennett et al., 1997). Loss of sialic acids results in decrease calcium ion channel sensitivity (Bennett et al., 1997), and perturbations of calcium homeostasis have been shown to impact DMD disease severity (Fong et al., 1990; Duan et al., 2021). These data suggest that decreased *MAN1A1* expression, like *KLOTHO* over-expression, may reduce sialic acids, which in turn may improve DMD pathology, as we observe via improved birefringence in *dmd-/-;man1a1* crispants (Figure 5). Future work investigating the role of *man1a1* in regulation of the immune response, calcium homeostasis, and/or myogenesis may shed light on how MAN1A1 influences DMD severity.

### *galntl6* as a modifier of DMD

GALNTL6 is a polypeptide N- acetylgalactosaminyltransferase involved in protein O-linked glycosylation. GALNTL6 is unable to initiate glycosylation, however once a peptide is modified with O-GalNAc, GALNTL6 can further glycosylate the peptide (Peng et al., 2010). Defects in O-linked glycosylation are linked to different congenital muscular dystrophies such as Walker-Warburg syndrome and muscle-eye-brain disease (Godfrey et al., 2007; Reily et al., 2019; Martin, 2010; Muntoni et al., 2007). Our data suggest that loss of *galntl6* leads to earlier disease onset in *dmd* mutant zebrafish (Figure 4) which is consistent with the observation that defects in O-linked glycosylation can lead to congenital muscular dystrophies. Other GWAS has shown *GALNTL6* coding sequence variants correlate with power (anaerobic) or endurance (aerobic) athletic performance (Díaz Ramírez et al., 2020). Single nuclei transcriptomic analysis of human muscle indicates that *GALNTL6* expression is higher in fast-twitch, oxidative muscle tissue (Nieves-Rodriguez et al., 2023). Thus, one plausible mechanism of GALNTL6 as a DMD modifier could be by altering muscle physiology or type. Shifting muscle metabolism towards an oxidative state, rather than glycolytic state, has been shown previously to be protective in DMD (Chakkalakal et al., 2004; Stupka et al., 2006), consistent with the observation that oxidative muscle is more resistant to Dystrophin loss than glycolytic muscle (Webster et al., 1988). Loss of *galntl6* in zebrafish increased disease onset but did not influence disease severity at the time point analyzed (4 dpf); future experiments will determine the effect of over-expressing *galntl6*, and human gene variants.

### *etaa1* as a modifier of DMD

ETAA1 is an activating protein for ATR which is involved in regulating replication stress and DNA damage response pathways. Mice deficient in *Etaa1* have defects in processes requiring in rapid cell proliferation (Miosge et al., 2017). Rapid cell proliferation of muscle stem cells (MuSCs) is a key hallmark in early stages of DMD; MuSCs are activated, differentiate, and proliferate to regenerate damaged muscle tissue whereby MuSCs fail to regenerate and exacerbate DMD severity (Filippelli & Chang, 2022; Kodippili & Rudnicki, 2023). There is conflicting evidence in the field regarding MuSC number and outcome of DMD severity. It has been shown that increasing the population of MuSCs improves DMD severity (Kodippili & Rudnicki, 2023) but conversely that MuSC depletion improves symptoms in both *mdx* and *sgcd* (a model of limb girdle MD caused by mutations in the delta-sarcoglycan gene) mice (Boyer et al., 2022). These data highlight the need to better understand the context- and temporal-dependent function of MuSCs in DMD pathology. Our data suggest that decreased expression of *ETAA1* ameliorates DMD severity at 4 dpf (Figure 6). Further experiments will interrogate the number, status, and function of MuSCs in *dmd-/-;etaa1a;etaa1b* crispants and the relevant controls.

### *adamts17* as a modifier of DMD

The ADAMTS family of enzymes are secreted metalloendopeptidases that regulate ECM formation and remodeling (Kelwick et al, 2015; Mead and Apte, 2018). Because the ADAMTS family encode secreted enzymes, they are appealing therapeutic targets, and drug development for some ADAMTS17 family members is underway for arthritis, neurodegenerative disease, and thrombocytopenia (Ferguson, 2025; Scully et al., 2017; van der Aar, et al., 2021; Bihlet et al., 2024; Schnitzer et al., 2023). The substrates of ADAMTS enzymes are mostly ECM components, however substrates of ADAMTS17 and ADAMTS19, two closely related sister enzymes, are not fully verified (Kelwick et al, 2015; Mead and Apte, 2018; Karoulias et al, 2020). As previously mentioned, we postulate that zebrafish Adamts17 performs the function that both ADAMTS17 and ADAMTS19 perform in humans. Interestingly, ADAMTS17 has been shown to bind, but not cleave, Fibrillin-2, an ECM component (Karoulias et al, 2020) and *adamts17* mutant mice have increased deposition of Fibrillin-2 in humeral distal growth plates (Oichi et al., 2019). The rs12657665 SNP that identified the lead candidate modifier gene *ADAMTS19* is also linked to *FBN2*, the gene that encodes Fibrillin-2 (Flanigan et al, 2023). It has been proposed that a regulatory network of Fibrillin (Fibrillin 1 and 2) microfibrils brings LTBPs and ADAMTS proteases together to regulate TGFβ signaling in the ECM (Karoulias et al, 2020) and we and others have shown that inhibition of TGFβ signaling ameliorates muscular dystrophies (Delaney et al, 2017; Fig. S2). In addition to (or in combination with) regulating TGFβ, Adamts17 may alter ECM composition or function, which has also been shown to modify DMD severity (Ceco and McNally, 2013; Burks and Chon, 2011). Thus, in our experiments, loss of *adamts17* function may delay disease onset, ameliorate DMD severity, and increase survivorship (Figure 7) by decreasing TGFβ signaling or altering ECM, and current experiments are focusing on characterizing ECM composition and TGFβ signaling responses to better understand the mechanism of action.

## Conclusions

We developed a CRISPR/Cas9 screening pipeline to identify modifiers of DMD. We showed that knockdown of homologs of known DMD modifiers *LTBP4* and *THBS1* affected DMD severity in zebrafish. We then validated that *galntl6, man1a1, etaa1a;etaa1b,* and *adamts17* are *bona fide* DMD modifiers. Further characterization of these modifiers, and additional human disease modifiers identified via GWAS, and the mechanisms by which they impact DMD pathology will identify the most promising for development of new DMD therapeutics.

## Author Contributions

G.T.L, K.R.R, J.C.B, O.M.C, and S.L.A performed experiments and analyzed data. All authors contributed intellectually and discussed the data and manuscript. G.T.L drafted the manuscript and all authors participated in the editing process.

## Acknowledgements

We thank the Ohio State Zebrafish facilities staff for excellent zebrafish care, Danielle Pvirre for technical assistance, and the entire Amacher lab for input and advice. We thank Drs. Kevin Flanigan, Veronica Vieland, and Robert Weiss and other members of the United Dystrophinopathy Project for advice and sharing GWAS data prior to publication. We thank Dr. James Jontes for sharing equipment, Dr. Thomas Gallagher for qPCR advice, Dr. Jared Talbot for imaging advice, Drs. Mayssa Mokalled and Dana Klatt Shaw for advice on CRISPR-based mutagenesis, Dr. Robert Weiss for contributions to Supplemental Figure 3, and Dr. John Postlethwait for discussions about ADAMTS17/19 family genes. We thank Drs. Veronica Vieland and Robert Weiss for reviewing manuscript drafts and providing constructive comments.

## Materials and Methods

### Animal Stocks and husbandry

Adult zebrafish strains (*Danio rerio*) were kept at 28.5°C on a 14h light/dark cycle and obtained by natural spawning or *in vitro* fertilization and were staged as previously described (Kimmel 1995). The *dmd^ta222a^* line has been described previously (Granato et al., 1996; M. Li et al., 2017) and has been maintained in our facility on the AB6 wild-type background for many generations.

### RNA extraction, cDNA transcription, and quantitative PCR

Whole embryos at 4, 7, and 10 dpf (n=15 per time point) were placed in Trizol, homogenized with a micro pestle for RNA extraction, and purified following standard procedures (Thermo Fisher). Total RNA (0.5 mg) was used for reverse transcription using random primers and Superscript IV reverse transcriptase (RT) according to the manufacturer’s instructions. Quantitative PCR (qPCR) was performed using PowerUp SYBER Green Master Mix (Thermo Fisher) and 4.5ul cDNA (diluted 1:50) in 20ul reactions, following manufacturer’s procedures. Negative controls lacking template were included for each primer set. For each time point, three biological replicates were performed, and transcript levels were normalized to *eef1a1a*. Cycle thresholds (C_t_) were determined using Bio-Rad CFX manager software. Changes in mRNA expression were calculated by ΔΔC_t_ = ΔC_t target_ - ΔC_t control_. Relative changes in mRNA expression levels are represented graphically as fold change, where relative mRNA fold change = 2^-ΔΔCt^ All qPCR primers are listed in Supplemental Table 1.

### DNA extraction and dmdta222a genotyping

To extract DNA, individual embryos and adult fin tissue were lysed in 30ul 1X ThermoPol Buffer (NEB) at 95°C for 10min, digested at 55°C for 1-4 hr using 25-50ug Proteinase K (ThermoFisher), followed by Proteinase K inactivation at 95°C for 10min. 1ul of DNA extract was used as template in a 20uL PCR reaction with Taq polymerase. For *dmd* genotyping, PCR was carried out with forward (5’- CATACCCAAGGTTTCAAAGCA -3’) and reverse (5’-TGCACTCGAGTGAAGCCACGTTTTT-3’) primers. PCR reactions were then digested with DraI (20 units) at 37°C for 30 minutes and analyzed on a 2% agarose gel stained with Gel Red (Biotium) to distinguish *dmd* wild-type (140 bp product) and *dmd^ta222a^* (110 bp product) alleles.

### CRISPR/Cas9 mutagenesis

CRISPR/Cas9 design and mutagenesis were performed as previously described (Shaw & Mokalled, 2021). crRNA guide RNA sequences were selected using either CHOPCHOP or IDT to target (and thus disrupt) essential protein domains of each gene of interest. Prior to CRISPR microinjection, adult *dmd+/-* zebrafish were screened via HRMA with each designated primer set to confirm a single cluster was detected via HRMA. If HRMA revealed more than one cluster, possibly indicating that SNPs in the interval around the target site were segregating in the population, we identified adult pairs that yielded a single HRMA cluster to simplify HRMA mutagenesis detection.

Lyophilized Alt-R tracrRNA and crRNA gRNAs (IDT, Cat# 1072534) were prepared as previously described (Shaw & Mokalled, 2021). In short, tracrRNA (100uM) and crRNA (100uM) were mixed (50uM), annealed by heating to 95°C and gradual cooling to 25°C, diluted to 25uM and stored at -20°C until use. Cas9 protein (QB3-Berkeley, Cas9-NLS Purified Protein [2.5mg]) was diluted to 25uM and stored at -80°C. For injection, sgRNA duplexes and Cas9 protein were mixed 1:1 in equal molar amounts with 1M KCl (0.2M final concentration) and Phenol Red (X% final concentration). Progeny obtained from intercrosses of *dmd* heterozygous adults were injected with 1nL of CRISPR/Cas9 solution at the one-cell stage and raised for phenotypic assays. All injected fish were genotyped via HRMA to identify whether the target region was mutagenized when compared to a subset of uninjected wild-type controls. All CRISPR guides are listed in Supplemental Table 2.

### High-resolution melt analysis

High-resolution melt analysis (HRMA) was conducted as previously described (Talbot & Amacher, 2014) to identify if mutagenesis had occurred at the target site. In short, primers were designed (Primer3) flanking the CRISPR target site. DNA was extracted from adult fin tissue or whole larvae as described above and 1ul was used as template in a 10ul HRMA reaction (BioRad 172-5112) in a CFX Duet Real-Time PCR System (#12016265) and analyzed using BioRad Precision Melt Analysis Software. Difference curves and melt peaks were analyzed to determine if mutagenesis had occurred (Fig. S1). All HRMA primers are listed in Supplemental Table 3.

### DMD onset determination

DMD is variable in its progression. In our experiments, *dmd* mutants begin showing birefringence anomalies at 2 dpf and all display birefringence anomalies by 4 dpf. To determine if modifier mutations impact DMD onset, we examined each fish daily to determine the first day at least one birefringence anomaly appears. Live fish were transferred to a 35mm glass bottom dish for birefringence scoring and those showing one or more birefringence anomaly (a DMD birefringence phenotype) were counted and moved to a new 100mm plate while the remaining fish showing no anomalies (a wild-type birefringence phenotype) were returned to the original plate. This process was repeated on 3 dpf and 4 dpf. Each day, fish scored for a DMD birefringence phenotype the previous day were rescored to confirm proper phenotyping (i.e. at 3 dpf, 2 dpf sorted for DMD phenotype were placed back into the glass bottom dish to confirm phenotyping). This assay identifies when one or more lesions first appear, however, it is a qualitative assay because a fish that is scored as showing a DMD phenotype could have just one anomaly (very mild) or multiple anomalies (severe). There is no correlation between day of onset (between day 2 and 4) and birefringence intensity quantification at 4 dpf (data not shown, R^2^ = .02). After scoring on 4 dpf, fish were fixed in 4% PFA overnight, rinsed 5 times for 10 minutes in PBST and processed for birefringence imaging (see below) and HRMA.

### Birefringence Imaging and Analysis

Birefringence was used to quantitatively assess dystrophy severity (Berger et al., 2012). Zebrafish were raised to 4 dpf, fixed overnight in 4% PFA, and rinsed with PBST 5 times for 10 minutes. A 35mm glass bottom dish was previously prepared for imaging by using a 1% agar solution and plastic mold (Megason, 2009) to create ‘inserts’ to hold fish for ease of imaging. Fish were individually placed into inserts for imaging. Using a Leica MZFLIII stereomicroscope equipped with polarizing filters, birefringence images were taken with a Zeiss AxioCam Hrc camera using AxioVision SE64 Rel 4.8 software. ImageJ was used to quantify mean gray values using the polygon selection tool to outline the entire trunk of the fish. Mean grey values are presented as a percentage of the average mean gray value of the WT siblings in their respective control group.

### Survivorship Assay

A survivorship assay was performed to determine candidate modifier mutation impacts of wild-type or *dmd* mutant fish lifespan. At 5 dpf, larval zebrafish were placed into 10cm dishes at a max density of 20 fish per dish and reared in a 28.5°C incubator with a 16 hour light:8 hour dark light cycle. Fish were fed paramecium once daily and received ∼75% water changes every other day. At lights on, fish were checked for survivorship. Between 5 and 12 dpf, dead fish were removed and placed into a single tube of an 8- well strip for DNA extraction and genotyping. After collecting dead fish on 12 dpf, all surviving fish were sacrificed and placed into individual tubes for DNA extraction and genotyping.

### ADAMTS19 Homology Comparisons

Protein sequences of human ADAMTS17 (ENST00000268070.9, Q8TE56-1) and ADAMTS19 (ENST00000274487.9, A0A1X7SBR9) and zebrafish Adamts17 (ENSDART00000113429.4, A0A8M9QHG7) were retrieved. Domain coordinates for the peptidase M12B, disintegrin, and TSP-type 1 domains were defined from InterProScan using the CDD and SMART databases. Domain percent identity and percent similarity using EMBOSS needle. The PLAC and spacer domains were not annotated on the CDD and SMART databases. Uniprot annotations were used for these domains and percent identity and similarity were calculated using TcoffeWS alignment tool.

### Statistical analyses

All statistical analyses were performed in GraphPad Prism 9 unless noted. Significance for all tests was set to p<0.05. Birefringence data were analyzed using a two-tailed Mann-Whitney U test. Survivorship data were analyzed for trend using a Log-rank test. DMD onset data were analyzed via Chi-Square test in Microsoft Excel. Expected distributions for Chi-Square analyses were calculated using trends observed in respective control *dmd* mutant groups.

### Power analysis

A priori power analysis were carried out using G*Power 3.1 (Faul et al., 2007). DMD onset: Chi-square analysis to detect a moderate effect size (w=.3) determined a sample size of 108 was necessary (DF = 2, Power =.8). Birefringence mean grey values: T-test analysis of birefringence data to detect a moderate effect size (d=.5) determined a sample size of 64 was necessary (two-tail, Power =.8, allocation ratio of 1).

## References

1. Adams JC, Lawler J. (2011). The thrombospondins. Cold Spring Harb Perspect Biol. 3(10):a009712.

2. Ahrens HE, Huettemeister J, Schmidt M, Kaether C, von Maltzahn J. (2018). Klotho expression is a prerequisite for proper muscle stem cell function and regeneration of skeletal muscle. Skelet Muscle. 8: 20.

3. Bass TE, Luzwick JW, Kavanaugh G, Carroll C, Dungrawala H, Glick GG, Feldkamp MD, Putney R, Chazin WJ, Cortez D. (2016). ETAA1 acts at stalled replication forks to maintain genome integrity. Nat Cell Biol. 18(11):1185–1195.

4. Bass TE, Cortez D. (2019). Quantitative phosphoproteomics reveals mitotic function of the ATR activator ETAA1. J Cell Biol. 218: 1235–1249.

5. Bassett, D. I., Bryson-Richardson, R. J., Daggett, D. F., Gautier, P., Keenan, D. G., & Currie, P. D. (2003). Dystrophin is required for the formation of stable muscle attachments in the zebrafish embryo. Development. 130(23):5851–5860.

6. Bek JW, Shochat C, De Clercq A, De Saffel H, Boel A, Metz J, Rodenburg F, Karasik D, Willaert A, Coucke PJ. (2021). Lrp5 Mutant and Crispant Zebrafish Faithfully Model Human Osteoporosis, Establishing the Zebrafish as a Platform for CRISPR-Based Functional Screening of Osteoporosis Candidate Genes. J Bone Miner Res. 36(9):1749–1764.

7. Bello L, Flanigan KM, Weiss RB; United Dystrophinopathy Project; Spitali P, Aartsma-Rus A, Muntoni F, Zaharieva I, Ferlini A, Mercuri E, Tuffery-Giraud S, Claustres M, Straub V, Lochmüller H, Barp A, Vianello S, Pegoraro E, Punetha J, Gordish-Dressman H, Giri M, McDonald CM, Hoffman EP; Cooperative International Neuromuscular Research Group. (2016). Association Study of Exon Variants in the NF-κB and TGFβ Pathways Identifies CD40 as a Modifier of Duchenne Muscular Dystrophy. Am J Hum Genet. 99(5):1163–1171.

8. Bello L, Piva L, Barp A, Taglia A, Picillo E, Vasco G, Pane M, Previtali SC, Torrente Y, Gazzerro E, Motta MC, Grieco GS, Napolitano S, Magri F, D’Amico A, Astrea G, Messina S, Sframeli M, Vita GL, Boffi P, Mongini T, Ferlini A, Gualandi F, Soraru’ G, Ermani M, Vita G, Battini R, Bertini E, Comi GP, Berardinelli A, Minetti C, Bruno C, Mercuri E, Politano L, Angelini C, Hoffman EP, Pegoraro E. (2012). Importance of SPP1 genotype as a covariate in clinical trials in Duchenne muscular dystrophy. Neurology. 79(2):159–62.

9. Bello, L., & Pegoraro, E. (2019). The “usual suspects”: Genes for inflammation, fibrosis, regeneration, and muscle strength modify duchenne muscular dystrophy. Journal of Clinical Medicine 8(5).

10. Bennett E, Urcan MS, Tinkle SS, Koszowski AG, Levinson SR. (1997). Contribution of sialic acid to the voltage dependence of sodium channel gating. A possible electrostatic mechanism. J Gen Physiol. 109: 327–343.

11. Berger J, Sztal T, Currie PD. (2012). Quantification of birefringence readily measures the level of muscle damage in zebrafish. Biochem Biophys Res Commun. 423(4):785–8.

12. Bez Batti Angulski, A., Hosny, N., Cohen, H., Martin, A. A., Hahn, D., Bauer, J., & Metzger, J. M. (2023). Duchenne muscular dystrophy: disease mechanism and therapeutic strategies. Frontiers in Physiology, 14, 1183101.

13. Bihlet AR, Balchen T, Goteti K, Sonne J, Ladel C, Karsdal MA, Ona V, Moreau F, Waterhouse R, Bay-Jensen AC, Guehring H. (2024). Safety, Tolerability, and Pharmacodynamics of the ADAMTS-5 Nanobody M6495: Two Phase 1, Single-Center, Double-Blind, Randomized, Placebo-Controlled Studies in Healthy Subjects and Patients With Osteoarthritis. ACR Open Rheumatol. 6(4):205–213.

14. Blazev R, Ashwood C, Abrahams JL, Chung LH, Francis D, Yang P, Watt KI, Qian H, Quaife-Ryan GA, Hudson JE, Gregorevic P, Thaysen-Andersen M, Parker BL. (2020). Integrated Glycoproteomics Identifies a Role of N-Glycosylation and Galectin-1 on Myogenesis and Muscle Development. Mol Cell Proteomics. 20:100030.

15. Boehler, J. F., Brown, K. J., Beatka, M., Gonzalez, J. P., Donisa Dreghici, R., Soustek-Kramer, M., McGonigle, S., Ganot, A., Palmer, T., Lowie, C., Chamberlain, J. S., Lawlor, M. W., & Morris, C. A. (2023). Clinical potential of microdystrophin as a surrogate endpoint. Neuromuscular Disorders, 33(1), 40–49.

16. Brunet FG, Fraser FW, Binder MJ, Smith AD, Kintakas C, Dancevic CM, Ward AC, McCulloch DR. (2015). The evolutionary conservation of the A Disintegrin-like and Metalloproteinase domain with Thrombospondin-1 motif metzincins across vertebrate species and their expression in teleost zebrafish. BMC Evol Biol.15:22.

17. Boyer, J. G., Huo, J., Han, S., Havens, J. R., Prasad, V., Lin, B. L., Kass, D. A., Song, T., Sadayappan, S., Khairallah, R. J., Ward, C. W., & Molkentin, J. D. (2022). Depletion of skeletal muscle satellite cells attenuates pathology in muscular dystrophy. Nature Communications, 13(1), 2940.

18. Bushby, K., Finkel, R., Birnkrant, D. J., Case, L. E., Clemens, P. R., Cripe, L., Kaul, A., Kinnett, K., McDonald, C., Pandya, S., Poysky, J., Shapiro, F., Tomezsko, J., & Constantin, C. (2010). Diagnosis and management of Duchenne muscular dystrophy, part 1: diagnosis, and pharmacological and psychosocial management. The Lancet Neurology, 9(1), 77–93.

19. Burks TN, Cohn RD. (2011). Role of TGF-β signaling in inherited and acquired myopathies. Skelet Muscle. 1(1):19.

20. Callahan JF, Burgess JL, Fornwald JA, Gaster LM, Harling JD, Harrington FP, Heer J, Kwon C, Lehr R, Mathur A, Olson BA, Weinstock J, Laping NJ. (2002). Identification of novel inhibitors of the transforming growth factor beta1 (TGF-beta1) type 1 receptor (ALK5). J. Med. Chem., 45, 999–1001.

21. Capote J., Kramerova I., Martinez L., Vetrone S., Barton E.R., Sweeney H.L., Miceli M.C. and Spencer M.J. (2016) Osteopontin ablation ameliorates muscular dystrophy by shifting macrophages to a pro-regenerative phenotype. J. Cell Biol. 213, 275–288.

22. Ceco E, McNally EM. (2013). Modifying muscular dystrophy through transforming growth factor-β. FEBS J. 280(17):4198–209.

23. Chakkalakal JV, Harrison MA, Carbonetto S, Chin E, Michel RN, Jasmin BJ. (2004). Stimulation of calcineurin signaling attenuates the dystrophic pathology in mdx mice. Hum Mol Genet. 13(4):379–88.

24. Crisafulli S, Sultana J, Fontana A, Salvo F, Messina S, Trifirò G. (2020). Global epidemiology of Duchenne muscular dystrophy: an updated systematic review and meta-analysis. Orphanet J Rare Dis. 5;15(1):141

25. Dahlem TJ, Hoshijima K, Jurynec MJ, Gunther D, Starker CG, Locke AS, Weis AM, Voytas DF, Grunwald DJ. (2012). Simple methods for generating and detecting locus-specific mutations induced with TALENs in the zebrafish genome. PLoS Genet. 8(8).

26. D’Ambrosio, E. S., & Mendell, J. R. (2023). Evolving Therapeutic Options for the Treatment of Duchenne Muscular Dystrophy. Neurotherapeutics : The Journal of the American Society for Experimental NeuroTherapeutics, 20(6), 1669–1681.

27. Debaenst, S., Jarayseh, T., Saffel, H., Bek, J. W., Boone, M., Josipovic, I., Kibleur, P., Kwon, R. Y., Coucke, P. J., Willaert, A. (2024) Crispant analysis in zebrafish as a tool for rapid functional screening of disease-causing genes for bone fragility. eLife. 13.

28. Delaney K, Kasprzycka P, Ciemerych MA, Zimowska M. (2017). The role of TGF-β1 during skeletal muscle regeneration. Cell Biol Int. 41(7):706–715.

29. Demonbreun, A. R., Fallon, K. S., Oosterbaan, C. C., Vaught, L. A., Reiser, N. L., Bogdanovic, E., Velez, M. P., Salamone, I. M., Page, P. G. T., Hadhazy, M., Quattrocelli, M., Barefield, D. Y., Wood, L. D., Gonzalez, J. P., Morris, C., & McNally, E. M. (2021). Anti-latent TGFβ binding protein 4 antibody improves muscle function and reduces muscle fibrosis in muscular dystrophy. Science Translational Medicine, 13(610).

30. Díaz Ramírez, J., Álvarez-Herms, J., Castañeda-Babarro, A., Larruskain, J., Ramírez de la Piscina, X., Borisov, O. V., Semenova, E. A., Kostryukova, E. S., Kulemin, N. A., Andryushchenko, O. N., Larin, A. K., Andryushchenko, L. B., Generozov, E. V., Ahmetov, I. I., & Odriozola, A. (2020). The GALNTL6 Gene rs558129 Polymorphism Is Associated With Power Performance. Journal of Strength and Conditioning Research. 34(11), 3031–3036.

31. DiMario JX. (2018). KLF10 Gene Expression Modulates Fibrosis in Dystrophic Skeletal Muscle. Am J Pathol. 188(5):1263–1275.

32. Duan, D., Goemans, N., Takeda, S., Mercuri, E., & Aartsma-Rus, A. (2021). Duchenne muscular dystrophy. Nature Reviews Disease Primers, 7(1), 13.

33. Duda T, Pertzev A, Ravichandran S, Sharma RK. (2018). Ca(2+)-Sensor Neurocalcin delta and Hormone ANF Modulate ANF-RGC Activity by Diverse Pathways: Role of the Signaling Helix Domain. Front Mol Neurosci. 11: 430.

34. Dumont NA, Wang YX, von Maltzahn J, Pasut A, Bentzinger CF, Brun CE, Rudnicki MA. (2015). Dystrophin expression in muscle stem cells regulates their polarity and asymmetric division. Nat Med. 21(12):1455–63

35. El-Brolosy, M. A., & Stainier, D. Y. R. (2017). Genetic compensation: A phenomenon in search of mechanisms. PLoS Genetics, 13(7).

36. Emery, AE. (1991) Population frequencies of inherited neuromuscular diseases—a world survey. Neuromuscul Disord. 1;1:19–29.

37. Esmail S, Manolson MF. (2021). Advances in understanding N-glycosylation structure, function, and regulation in health and disease. Eur J Cell Biol. 100(7-8):151186.

38. Evans, D. R., Green, J. S., Fahiminiya, S., Majewski, J., Fernandez, B. A., Deardorff, M. A., Johnson, G. J., Whelan, J. H., Hubmacher, D., Apte, S. S., Boycott, K., Bulman, D., Dyment, D., McKenzie, A., Brudno, M., & Woods, M. O. (2020). A novel pathogenic missense ADAMTS17 variant that impairs secretion causes Weill-Marchesani Syndrome with variably dysmorphic hand features. Scientific Reports. 10(1), 10827.

39. Farr GH 3rd, Morris M, Gomez A, Pham T, Kilroy E, Parker EU, Said S, Henry C, Maves L. A. (2020). novel chemical-combination screen in zebrafish identifies epigenetic small molecule candidates for the treatment of Duchenne muscular dystrophy. Skelet Muscle. 10(1):29.

40. Faul, F., Erdfelder, E., Lang, A.-G., & Buchner, A. (2007). G*Power 3: A flexible statistical power analysis program for the social, behavioral, and biomedical sciences. Behavior Research Methods. 39, 175–191.

41. Feige P, Brun CE, Ritso M, Rudnicki MA. (2018). Orienting muscle stem cells for regeneration in homeostasis, aging, and disease. Cell Stem Cell. 23:653–64.

42. Ferguson, MRSB MRSC D. (2025). Integrative Design of ADAMTS Partial Agonists within a Multi-Hallmark Therapeutic Framework: The Beginning of Paving the Way Toward a Functional Cure for Neurodegenerative Disorders. ChemRxiv. doi:10.26434/chemrxiv-2025-4n2nb-v3

43. Filippelli, R. L., & Chang, N. C. (2022). Empowering Muscle Stem Cells for the Treatment of Duchenne Muscular Dystrophy. Cells Tissues Organs, 211(6), 641– 654.

44. Flanigan, K. M., Waldrop, M. A., Martin, P. T., Alles, R., Dunn, D. M., Alfano, L. N., Simmons, T. R., Moore-Clingenpeel, M., Burian, J., Seok, S. C., Weiss, R. B., & Vieland, V. J. (2023). A genome-wide association analysis of loss of ambulation in dystrophinopathy patients suggests multiple candidate modifiers of disease severity. European Journal of Human Genetics, 31(6), 663–673.

45. Flanigan, K. M., Ceco, E., Lamar, K. M., Kaminoh, Y., Dunn, D. M., Mendell, J. R., King, W. M., Pestronk, A., Florence, J. M., Mathews, K. D., Finkel, R. S., Swoboda, K. J., Gappmaier, E., Howard, M. T., Day, J. W., McDonald, C., McNally, E. M., Weiss, R. B., & United Dystrophinopathy Project. (2013). LTBP4 genotype predicts age of ambulatory loss in Duchenne muscular dystrophy. Annals of neurology, 73(4), 481–488.

46. Fong PY, Turner PR, Denetclaw WF, Steinhardt RA. (1990). Increased activity of calcium leak channels in myotubes of Duchenne human and mdx mouse origin. Science. 250: 673–676.

47. Geijtenbeek TB, Gringhuis SI. (2009). Signalling through C-type lectin receptors: shaping immune responses. Nat Rev Immunol. 9(7):465–79.

48. Granados A, Zamperoni M, Rapone R, Moulin M, Boyarchuk E, Bouyioukos C, Del Maestro L, Joliot V, Negroni E, Mohamed M, Piquet S, Bigot A, Le Grand F, Albini S, Ait-Si-Ali S. (2024). SETDB1 modulates the TGFβ response in Duchenne muscular dystrophy myotubes. Sci Adv. 8042.

49. Guglieri, M., Bushby, K., McDermott, M. P., Hart, K. A., Tawil, R., Martens, W. B., Herr, B. E., McColl, E., Speed, C., Wilkinson, J., Kirschner, J., King, W. M., Eagle, M., Brown, M. W., Willis, T., Griggs, R. C., Straub, V., van Ruiten, H., Childs, A.-M., Chang, T. (2022a). Effect of Different Corticosteroid Dosing Regimens on Clinical Outcomes in Boys With Duchenne Muscular Dystrophy. JAMA, 327(15), 1456.

50. Guglieri, M., Clemens, P. R., Perlman, S. J., Smith, E. C., Horrocks, I., Finkel, R. S., Mah, J. K., Deconinck, N., Goemans, N., Haberlova, J., Straub, V., Mengle- Gaw, L. J., Schwartz, B. D., Harper, A. D., Shieh, P. B., De Waele, L., Castro, D., Yang, M. L., Ryan, M. M., Hoffman, E. P. (2022b). Efficacy and Safety of Vamorolone vs Placebo and Prednisone Among Boys With Duchenne Muscular Dystrophy. JAMA Neurology, 79(10), 1005.

51. Heydemann A, Ceco E, Lim JE, Hadhazy M, Ryder P, Moran JL, Beier DR, Palmer AA, McNally EM. (2009). Latent TGF-beta-binding protein 4 modifies muscular dystrophy in mice. J Clin Invest. 119(12):3703–12.

52. Hinz B. (2015). The extracellular matrix and transforming growth factor-β1: Tale of a strained relationship. Matrix Biol. 47:54–65.

53. Hogarth MW, Houweling PJ, Thomas KC, Gordish-Dressman H, Bello L; Cooperative international neuromuscular research group (CINRG); Pegoraro E, Hoffman EP, Head SI, North Kn. (2017) Evidence for ACTN3 as a genetic modifier of Duchenne muscular dystrophy. Nat Commun. 31;8:14143.

54. Icer MA, Gezmen-Karadag M. (2018). The multiple functions and mechanisms of osteopontin. Clin Biochem. 59:17–24.

55. Inman GJ, Nicolas FJ, Callahan JF, Harling JD, Gaster LM, Reith AB, Lapin NJ, Hill CS. (2002). SB-431542 is a potent and specific inhibitor of transforming growth factor-beta superfamily type I activin receptor-like kinase (ALK) receptors ALK4, ALK5, and ALK7. 62, 65–74.

56. Ishitobi M, Haginova K, Zhao YJ, Ohnuma A, Minato J, Yanagisawa T, Tanabu M, Kikuchi M, Iinuma K. (2000) Elevated plasma levels of transforming growth factor β1 in patients with muscular dystrophy. Neuroreport. 11, 4033–4035.

57. Ismaeel A, Kim JS, Kirk JS, Smith RS, Bohannon WT, Koutakis P. (2019). Role of Transforming Growth Factor-β in Skeletal Muscle Fibrosis: A Review. Int J Mol Sci. 20(10):2446.

58. Jin, Z.-C., Kitajima, T., Dong, W., Huang, Y.-F., Ren, W.-W., Guan, F., Chiba, Y., Gao, X.-D., & Fujita, M. (2018). Genetic disruption of multiple α1,2-mannosidases generates mammalian cells producing recombinant proteins with high-mannose– type N-glycans. Journal of Biological Chemistry. 293(15), 5572–5584.

59. Johnson NM, Farr GH 3rd, Maves L. (2013). The HDAC Inhibitor TSA Ameliorates a Zebrafish Model of Duchenne Muscular Dystrophy. PLoS Curr. 17;5.

60. Juban G, Saclier M, Yacoub-Youssef H, Kernou A, Arnold L, Boisson C, Larbi SB, Magnan M, Cuvellier S, Theret M, Petrof BJ, Desguerre I, Gondin J, Mounier R, Chazaud B. (2018) AMPK Activation Regulates LTBP4-Dependent TGF- β1 Secretion by Pro-inflammatory Macrophages and Controls Fibrosis in Duchenne Muscular Dystrophy. Cell Reports. 25(8)2163–2176.

61. Karlsson M, Zhang C, Méar L, Zhong W, Digre A, Katona B, Sjöstedt E, Butler L, Odeberg J, Dusart P, Edfors F, Oksvold P, von Feilitzen K, Zwahlen M, Arif M, Altay O, Li X, Ozcan M, Mardinoglu A, Fagerberg L, Mulder J, Luo Y, Ponten F, Uhlén M, Lindskog C. (2021). A single-cell type transcriptomics map of human tissues. Sci Adv. 7(31).

62. Karoulias, S. Z., Taye, N., Stanley, S., & Hubmacher, D. (2020). The ADAMTS/Fibrillin Connection: Insights into the Biological Functions of ADAMTS10 and ADAMTS17 and Their Respective Sister Proteases. Biomolecules, 10(4).

63. Kawahara G, Karpf JA, Myers JA, Alexander MS, Guyon JR, Kunkel LM. (2011). Drug screening in a zebrafish model of Duchenne muscular dystrophy. Proc Natl Acad Sci U S A. 108(13):5331–6.

64. Kawahara G, Kunkel LM. (2013). Zebrafish based small molecule screens for novel DMD drugs. Drug Discov Today Technol. 10(1):e91–6.

65. Kemaladewi DU, Pasteuning S, van der Meulen JW, van Heiningen SH, van Ommen GJ, Ten Dijke P, Aartsma-Rus A, ’t Hoen PA, Hoogaars WM. (2014). Targeting TGF-β Signaling by Antisense Oligonucleotide-mediated Knockdown of TGF-β Type I Receptor. Mol Ther Nucleic Acids. 3(4):e156.

66. Kodippili, K., & Rudnicki, M. A. (2023). Satellite cell contribution to disease pathology in Duchenne muscular dystrophy. Frontiers in Physiology. 14, 1180980.

67. Kramerova I, Kumagai-Cresse C, Ermolova N, Mokhonova E, Marinov M, Capote J, Becerra D, Quattrocelli M, Crosbie RH, Welch E, McNally EM, Spencer MJ. (2019). Spp1 (osteopontin) promotes TGFβ processing in fibroblasts of dystrophin-deficient muscles through matrix metalloproteinases. Hum Mol Genet. 28(20):3431–3442.

68. Kroll, F., Powell, G. T., Ghosh, M., Gestri, G., Antinucci, P., Hearn, T. J., Tunbak, H., Lim, S., Dennis, H. W., Fernandez, J. M., Whitmore, D., Dreosti, E., Wilson, S. W., Hoffman, E. J., & Rihel, J. (2021). A simple and effective f0 knockout method for rapid screening of behaviour and other complex phenotypes. ELife, 10, 1–34.

69. Kurosu H, Yamamoto M, Clark JD, Pastor JV, Nandi A, Gurnani P, McGuinness OP, Chikuda H, Yamaguchi M, Kawaguchi H, Shimomura I, Takayama Y, Herz J, Kahn CR, Rosenblatt KP, Kuro-o M. (2005). Suppression of aging in mice by the hormone Klotho. Science. 309(5742):1829–33.

70. Legler, K., Rosprim, R., Karius, T., Eylmann, K., Rossberg, M., Wirtz, R. M., Müller, V., Witzel, I., Schmalfeldt, B., Milde-Langosch, K., & Oliveira-Ferrer, L. (2018). Reduced mannosidase MAN1A1 expression leads to aberrant N-glycosylation and impaired survival in breast cancer. British Journal of Cancer. 118(6), 847–856.

71. Lund SA, Giachelli CM, Scatena M. (2009). The role of osteopontin in inflammatory processes. J Cell Commun Signal. 3(3-4):311–22.

72. Matsuo, M. (2021). Antisense Oligonucleotide-Mediated Exon-skipping Therapies: Precision Medicine Spreading from Duchenne Muscular Dystrophy. JMA Journal, 4(3), 232–240.

73. Massadeh S, Alhashem A, van de Laar IMBH, Alhabshan F, Ordonez N, Alawbathani S, Khan S, Kabbani MS, Chaikhouni F, Sheereen A, Almohammed I, Alghamdi B, Frohn-Mulder I, Ahmad S, Beetz C, Bauer P, Wessels MW, Alaamery M, Bertoli-Avella AM. (2020). ADAMTS19-associated heart valve defects: Novel genetic variants consolidating a recognizable cardiac phenotype. Clin Genet. 98(1):56–63.

74. Massagué J. (2012). TGFβ signalling in context. Nat Rev Mol Cell Biol. 13(10):616–30.

75. Massagué J, Sheppard D. (2023). TGF-β signaling in health and disease. Cell. 186(19):4007–4037.

76. Mázala DA, Novak JS, Hogarth MW, Nearing M, Adusumalli P, Tully CB, Habib NF, Gordish-Dressman H, Chen YW, Jaiswal JK, Partridge TA. (2020). TGF-β- driven muscle degeneration and failed regeneration underlie disease onset in a DMD mouse model. JCI Insight. 5(6):e135703.

77. Megason SG. (2009). In toto imaging of embryogenesis with confocal time-lapse microscopy. Methods Mol Biol. 546:317–32.

78. Mendell JR, Campbell K, Rodino-Klapac L, Sahenk Z, Shilling C, Lewis S, Bowles D, Gray S, Li C, Galloway G, Malik V, Coley B, Clark KR, Li J, Xiao X, Samulski J, McPhee SW, Samulski RJ, Walker CM. (2010). Dystrophin immunity in Duchenne’s muscular dystrophy. N Engl J Med. 363(15):1429–37.

79. Mendell JR, Connolly AM, Lehman KJ, Griffin DA, Khan SZ, Dharia SD, Quintana-Gallardo L, Rodino-Klapac LR. (2022). Testing preexisting antibodies prior to AAV gene transfer therapy: rationale, lessons and future considerations. Mol Ther Methods Clin Dev. 25:74–83.

80. Mendell JR, Shilling C, Leslie ND, Flanigan KM, al-Dahhak R, Gastier-Foster K, Keile, Dunn DM, Duval B, Aoyagi A, Hamil C, Mahoud M, Roush K, Bird L, Rankin C, Lily H, Street N, Chandrasekar R, Weiss RB. (2012). Evidence-based path to newborn screening for Duchenne muscular dystrophy. Annal of neurology. 71(3), 304–313

81. Miosge, L. A., Sontani, Y., Chuah, A., Horikawa, K., Russell, T. A., Mei, Y., Wagle, M. V., Howard, D. R., Enders, A., Tscharke, D. C., Goodnow, C. C., & Parish, I. A. (2017). Systems-guided forward genetic screen reveals a critical role of the replication stress response protein ETAA1 in T cell clonal expansion. Proceedings of the National Academy of Sciences, 114(26).

82. Muntoni, F., Brockington, M., Godfrey, C., Ackroyd, M., Robb, S., Manzur, A., Kinali, M., Mercuri, E., Kaluarachchi, M., Feng, L., Jimenez-Mallebrera, C., Clement, E., Torelli, S., Sewry, C. A., & Brown, S. C. (2007). Muscular dystrophies due to defective glycosylation of dystroglycan. Acta Myologica : Myopathies and Cardiomyopathies : Official Journal of the Mediterranean Society of Myology, 26(3), 129–135.

83. Nieves-Rodriguez S, Barthélémy F, Woods JD, Douine ED, Wang RT, Scripture-Adams DD, Chesmore KN, Galasso F, Miceli MC, Nelson SF. (2023). Transcriptomic analysis of paired healthy human skeletal muscles to identify modulators of disease severity in DMD. Front Genet. 14:1216066.

84. Oichi T, Taniguchi Y, Soma K, Oshima Y, Yano F, Mori Y, Chijimatsu R, Kim-Kaneyama JR, Tanaka S, Saito T. Adamts17 is involved in skeletogenesis through modulation of BMP-Smad1/5/8 pathway. (2019). *Cell Mol Life Sci*. 76(23):4795-4809.

85. Parvez S, Brandt ZJ, Peterson RT. (2023). Large-scale F0 CRISPR screens in vivo using MIC-Drop. Nat Protoc. 18(6):1841–1865.

86. Pegoraro E, Hoffman EP, Piva L, Gavassini BF, Cagnin S, Ermani M, Bello L, Soraru G, Pacchioni B, Bonifati MD, Lanfranchi G, Angelini C, Kesari A, Lee I, Gordish-Dressman H, Devaney JM, McDonald CM; Cooperative International Neuromuscular Research Group. (2011) SPP1 genotype is a determinant of disease severity in Duchenne muscular dystrophy. Neurology. 18;76(3):219–26.

87. Peng, C., Togayachi, A., Kwon, Y.-D., Xie, C., Wu, G., Zou, X., Sato, T., Ito, H., Tachibana, K., Kubota, T., Noce, T., Narimatsu, H., & Zhang, Y. (2010). Identification of a novel human UDP-GalNAc transferase with unique catalytic activity and expression profile. Biochemical and Biophysical Research Communications, 402(4), 680–686.

88. Pinho SS, Alves I, Gaifem J, Rabinovich GA. (2023). Immune regulatory networks coordinated by glycans and glycan-binding proteins in autoimmunity and infection. Cell Mol Immunol. 20(10):1101–1113.

89. Postlethwait JH, Yan YL, Gates MA, Horne S, Amores A, Brownlie A, Donovan A, Egan ES, Force A, Gong Z, Goutel C, Fritz A, Kelsh R, Knapik E, Liao E, Paw B, Ransom D, Singer A, Thomson M, Abduljabbar TS, Yelick P, Beier D, Joly JS, Larhammar D, Rosa F, Westerfield M, Zon LI, Johnson SL, Talbot WS. (1998). Vertebrate genome evolution and the zebrafish gene map. Nat Genet. 18(4):345–9.

90. Raman J, Guan Y, Perrine CL, Gerken TA, Tabak LA. (2012). UDP-N-acetyl-alpha-D-galactosamine:polypeptide N-acetylgalactosaminyltransferases: completion of the family tree. Gylcobiology. 22: 768–777.

91. Reily, C., Stewart, T. J., Renfrow, M. B., & Novak, J. (2019). Glycosylation in health and disease. Nature Reviews Nephrology, 15(6), 346–366.

92. Riessland M, Kaczmarek A, Schneider S, Swoboda KJ, Löhr H, Bradler C, Grysko V, Dimitriadi M, Hosseinibarkooie S, Torres-Benito L, Peters M, Upadhyay A, Biglari N, Kröber S, Hölker I, Garbes L, Gilissen C, Hoischen A, Nürnberg G, Nürnberg P, Walter M, Rigo F, Bennett CF, Kye MJ, Hart AC, Hammerschmidt M, Kloppenburg P, Wirth B. (2017). Neurocalcin Delta Suppression Protects against Spinal Muscular Atrophy in Humans and across Species by Restoring Impaired Endocytosis. Am J Hum Genet. 100(2):297–315.

93. Samarut E, Lissouba A, Drapeau P. (2016). A simplified method for identifying early CRISPR-induced indels in zebrafish embryos using High Resolution Melting analysis. BMC Genomics. 17, 547.

94. Schnitzer T, Pueyo M, Deckx H, van der Aar E, Bernard K, Hatch S, van der Stoep M, Grankov S, Phung D, Imbert O, Chimits D, Muller K, Hochberg MC, Bliddal H, Wirth W, Eckstein F, Conaghan PG. (2023). Evaluation of S201086/GLPG1972, an ADAMTS-5 inhibitor, for the treatment of knee osteoarthritis in ROCCELLA: a phase 2 randomized clinical trial. Osteoarthritis Cartilage. 31(7):985–994.

95. Scully M, Knöbl P, Kentouche K, Rice L, Windyga J, Schneppenheim R, Kremer Hovinga JA, Kajiwara M, Fujimura Y, Maggiore C, Doralt J, Hibbard C, Martell L, Ewenstein B. (2017). Recombinant ADAMTS-13: first-in-human pharmacokinetics and safety in congenital thrombotic thrombocytopenic purpura. Blood. 130(19):2055–2063.

96. Shaw, D. K., & Mokalled, M. H. (2021). Efficient CRISPR/Cas9 mutagenesis for neurobehavioral screening in adult zebrafish. G3: Genes, Genomes, Genetics, 11(8).

97. Song Y, Tao S, Liu Y, Long L, Yang H, Li Q, Liang J, Li X, Lu Y, Zhu H. (2017) Expression levels of TGF-beta1 and CTGF are associated with the severity of Duchenne muscular dystrophy. Exp. Ther. Med. 13, 1209–1214.

98. Spitali P, Zaharieva I, Bohringer S, Hiller M, Chaouch A, Roos A, Scotton C, Claustres M, Bello L, McDonald CM, Hoffman EP; CINRG Investigators; Koeks Z, Eka Suchiman H, Cirak S, Scoto M, Reza M, ’t Hoen PAC, Niks EH, Tuffery-Giraud S, Lochmüller H, Ferlini A, Muntoni F, Aartsma-Rus A. (2020). TCTEX1D1 is a genetic modifier of disease progression in Duchenne muscular dystrophy. Eur J Hum Genet. 28(6):815–825.

99. Spittau B, Krieglstein K. (2012). Klf10 and Klf11 as mediators of TGF-beta superfamily signaling. Cell Tissue Res. 347: 65–72.

100. Stupka N, Plant DR, Schertzer JD, Emerson TM, Bassel-Duby R, Olson EN, Lynch GS. (2006). Activated calcineurin ameliorates contraction-induced injury to skeletal muscles of mdx dystrophic mice. J Physiol. 575(Pt 2):645–56.

101. Subramaniam M, Harris SA, Oursler MJ, Rasmussen K, Riggs BL, Spelsberg TC. (1995). Identification of a novel TGF-beta-regulated gene encoding a putative zinc finger protein in human osteoblasts. Nucleic Acids Res. 23:4907–4912.

102. Talbot, J. C., & Amacher, S. L. (2014). A Streamlined CRISPR Pipeline to Reliably Generate Zebrafish Frameshifting Alleles. Zebrafish. 11(6), 583–585.

103. Tidball, J. G., Welc, S. S., & Wehling-Henricks, M. (2018). Immunobiology of Inherited Muscular Dystrophies. In Comprehensive Physiology, 8(4) 1313–1356.

104. Todorovic V, Rifkin DB. (2012). LTBPs, more than just an escort service. J Cell Biochem. 113(2):410–8.

105. Trubiroha A, Gillotay P, Giusti N, Gacquer D, Libert F, Lefort A, Haerlingen B, De Deken X, Opitz R, Costagliola S. (2018). A Rapid CRISPR/Cas-based Mutagenesis Assay in Zebrafish for Identification of Genes Involved in Thyroid Morphogenesis and Function. Sci Rep. 8(1):5647

106. Uhlén M, Fagerberg L, Hallström BM, Lindskog C, Oksvold P, Mardinoglu A, Sivertsson Å, Kampf C, Sjöstedt E, Asplund A, Olsson I, Edlund K, Lundberg E, Navani S, Szigyarto CA, Odeberg J, Djureinovic D, Takanen JO, Hober S, Alm T, Edqvist PH, Berling H, Tegel H, Mulder J, Rockberg J, Nilsson P, Schwenk JM, Hamsten M, von Feilitzen K, Forsberg M, Persson L, Johansson F, Zwahlen M, von Heijne G, Nielsen J, Pontén F. (2015). Tissue-based map of the human proteome. Science. 347(6220):1260419.

107. van der Aar E, Deckx H, Dupont S, Fieuw A, Delage S, Larsson S, Struglics A, Lohmander LS, Lalande A, Leroux E, Amantini D, Passier P. (2022). Safety, Pharmacokinetics, and Pharmacodynamics of the ADAMTS-5 Inhibitor GLPG1972/S201086 in Healthy Volunteers and Participants With Osteoarthritis of the Knee or Hip. Clin Pharmacol Drug Dev. 11(1):112–122.

108. Varki A. (2017). Biological roles of glycans. Glycobiology. 27(1):3–49.

109. Vetrone S.A., Montecino-Rodriguez E., Kudryashova E., Kramerova I., Hoffman E.P., Liu S.D., Miceli M.C. and Spencer M.J. (2009) Osteopontin promotes fibrosis in dystrophic mouse muscle by modulating immune cell subsets and intramuscular TGF-beta. J. Clin. Invest., 119, 1583–1594.

110. Vieira NM, Elvers I, Alexander MS, Moreira YB, Eran A, Gomes JP, Marshall JL, Karlsson EK, Verjovski-Almeida S, Lindblad-Toh K, Kunkel LM, Zatz M. (2015). Jagged 1 Rescues the Duchenne Muscular Dystrophy Phenotype. Cell. 163(5):1204–1213.

111. Webster, C., Silberstein, L., Hays, A. P., & Blau, H. M. (1988). Fast muscle fibers are preferentially affected in Duchenne muscular dystrophy. Cell. 52(4), 503–513.

112. Wehling-Henricks M, Li Z, Lindsey C, Wang Y, Welc SS, Ramos JN, Khanlou N, Kuro-O M, Tidball JG. (2016). Klotho gene silencing promotes pathology in the mdx mouse model of Duchenne muscular dystrophy. Hum Mol Genet. 25(12):2465–2482.

113. Weiss, R. B., Vieland, V. J., Dunn, D. M., Kaminoh, Y., & Flanigan, K. M. (2018). Long-range genomic regulators of THBS1 and LTBP4 modify disease severity in duchenne muscular dystrophy. Annals of Neurology, 84(2), 234–245.

114. Welc SS, Wehling-Henricks M, Kuro OM, Thomas KA, Tidball JG. (2020). Modulation of Klotho expression in injured muscle perturbs Wnt signalling and influences the rate of muscle growth. Exp Physiol. 105:132–147.

115. Widrick JJ, Kawahara G, Alexander MS, Beggs AH, Kunkel LM. (2019) Discovery of Novel Therapeutics for Muscular Dystrophies using Zebrafish Phenotypic Screens. J Neuromuscul Dis. 6(3):271–287.

116. Wu RS, Lam II, Clay H, Duong DN, Deo RC, Coughlin SR. (2018). A Rapid Method for Directed Gene Knockout for Screening in G0 Zebrafish. Dev Cell. 46(1):112–125.e4.

117. Wu, R., Song, Y., Wu, S., & Chen, Y. (2022). Promising therapeutic approaches of utrophin replacing dystrophin in the treatment of Duchenne muscular dystrophy. Fundamental Research, 2(6), 885–893.

118. Yamashita K, Suzuki A, Satoh Y, Ide M, Amano Y, Masuda-Hirata M, Hayashi YK, Hamada K, Ogata K, Ohno S. (2010). The 8th and 9th tandem spectrin-like repeats of utrophin cooperatively form a functional unit to interact with polarity- regulating kinase PAR-1b. Biochem Biophys Res Commun. 391(1):812–7.

119. Zhang Y, Hartmann HA, Satin J. (1999). Glycosylation influences voltage-dependent gating of cardiac and skeletal muscle sodium channels. J Membr Biol. 171: 195–207.

120. Zhao Y, Huang Z, Gao L, Ma H, Chang R. (2024). Osteopontin/SPP1: a potential mediator between immune cells and vascular calcification. Front Immunol. 15:1395596.

