## Supplemental Data for "Validation of Duchenne muscular dystrophy candidate modifiers using a CRISPR-Cas9-based approach in zebrafish"

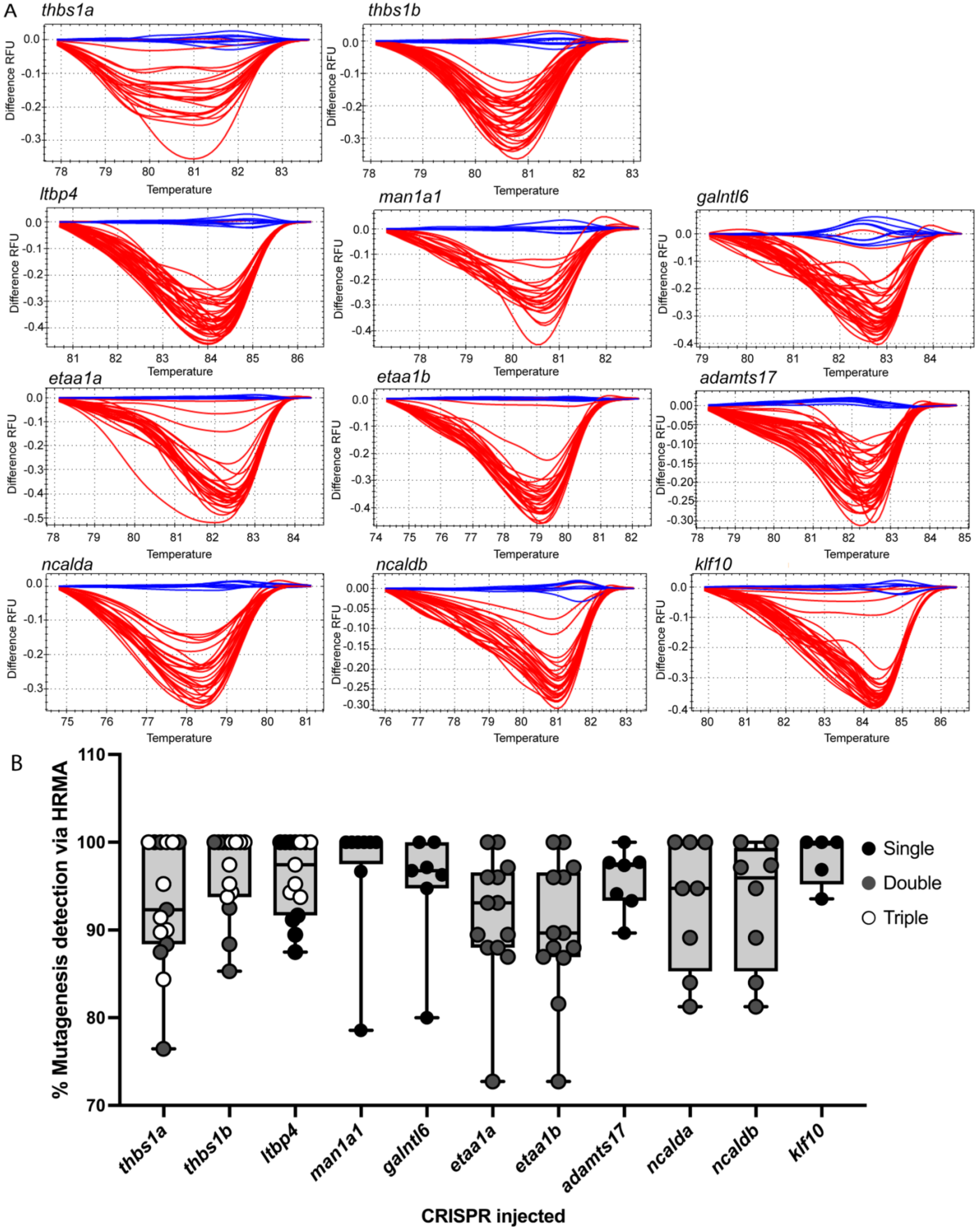

A
thbs1a
thbs1b
ltbp4
man1a1
galntl6
etaa1a
etaa1b
adamts17
ncalda
ncaldb
klf10
Difference RFU
Temperature
B
% Mutagenesis detection via HRMA
CRISPR injected
Single
Double
Triple

**Supplemental Figure 1: High resolution melt analysis demonstrates high-efficiency mutagenesis events in CRISPR/Cas9-microinjected embryos.** Following microinjection and embryo processing, all embryos were genotyped via HRMA. (A) Representative HRMA for each gene targeted. To compare differences in melting temperature between injected and uninjected fish, the difference in relative fluorescent units (RFU, y-axis) is calculated for each injected fish (red) versus temperature (x-axis), with uninjected fish (blue) set as the reference cluster. Crispants whose difference curves failed to deflect from the uninjected fish represented a small fraction (7.3%) of all injected fish and were omitted from subsequent analyses, including calculating of average mutagenesis percentage. (B) Box plot showing the average mutagenesis percentage in both WT and *dmd* mutants for each experiment conducted, per crispant condition. Data points are color coded based on number of CRISPRs injected in each experiment [one CRISPR (black), two CRISPRs (grey), three CRISPRs (white] and show that multiplexing CRISPRs does not appear to affect efficacy, as has been previously shown by others (Kroll et al., 2021; Wu et al., 2018).

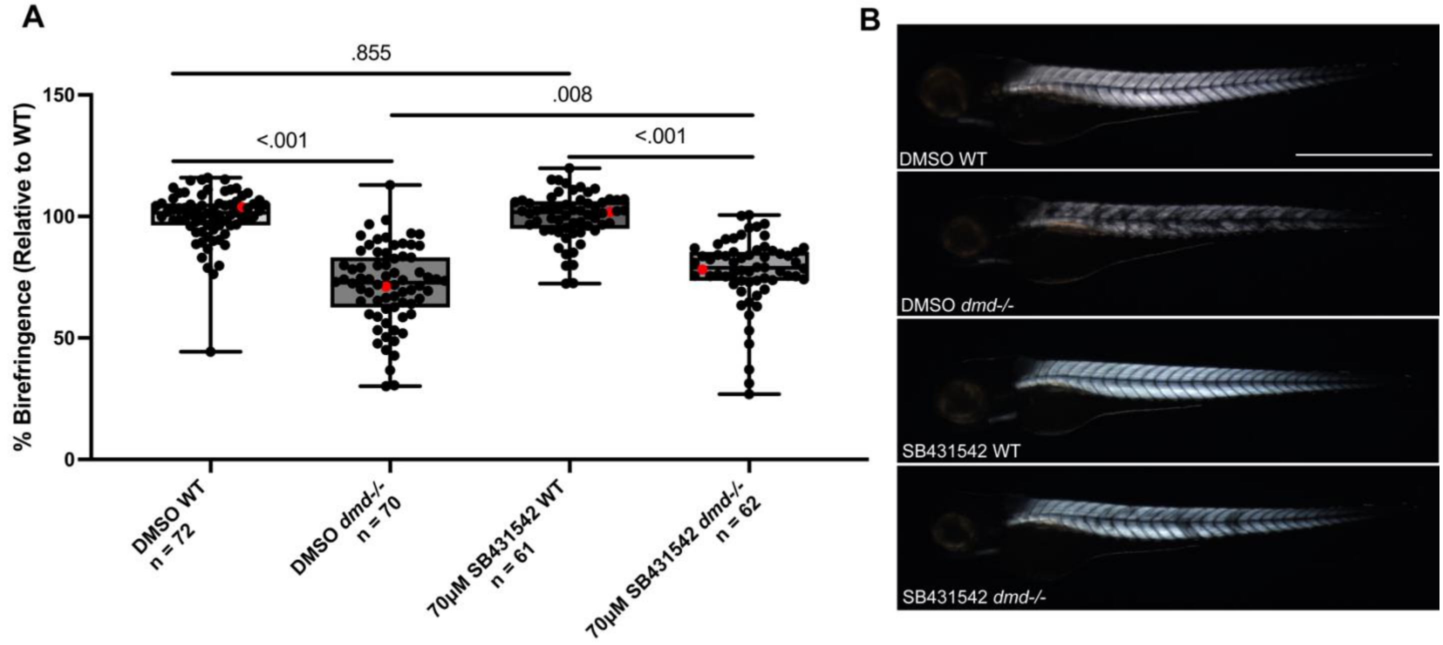


**Supplemental Figure 2: Treatment with SB431542 increases birefringence intensity.** Embryos from a *dmd+/-* intercross were raised to 24hpf and treated with DMSO or 70uM SB431542 until 48hpf, rinsed thoroughly, and raised to 4 dpf. (A) Box plots show overall birefringence intensities at 4 dpf. Mann-Whitney U-Tests carried out for comparisons of overall birefringence intensities. Red dots denote representative images shown in panel B. DMSO control and SB431542 treatment experiments were performed in biological triplicate. The number of fish (n) for each treatment condition is indicated. (B) Representative birefringence images for each treatment condition at 4 dpf. Scale bar denotes 1mm.

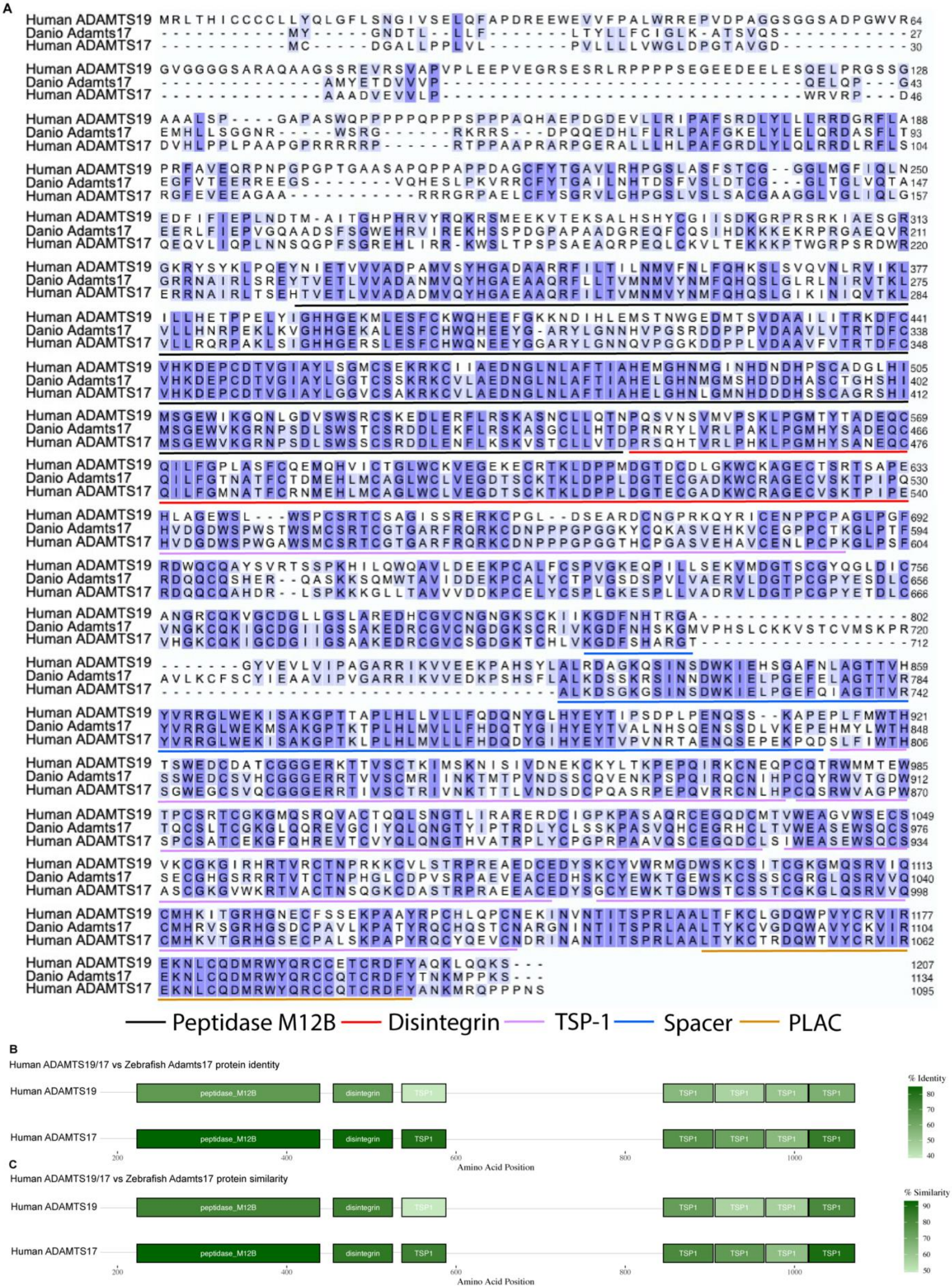
A
Human ADAMTS19
Danio Adamts17
Human ADAMTS17
Peptidase M12B
Disintegrin
TSP-1
Spacer
PLAC
B
Human ADAMTS19/17 vs Zebrafish Adamts17 protein identity
% Identity
C
Human ADAMTS19/17 vs Zebrafish Adamts17 protein similarity
% Similarity
peptidase_M12B
disintegrin
TSP1
Amino Acid Position

**Supplemental Figure 3. Zebrafish Adamts17 is similar to human ADAMTS19 and ADAMTS17.** The protein sequences Zebrafish Adamts17 (ENSDART00000113429.4, A0A8M9QHG7), Human ADAMTS19 (ENST00000274487.9, A0A1X7SBR9), and Human ADAMTS17 (ENST00000268070.9, Q8TE56-1) were compared. (A) Sequence identity and (B) Sequence similarity when comparing human ADAMTS19 and 17 to zebrafish Adamts17.

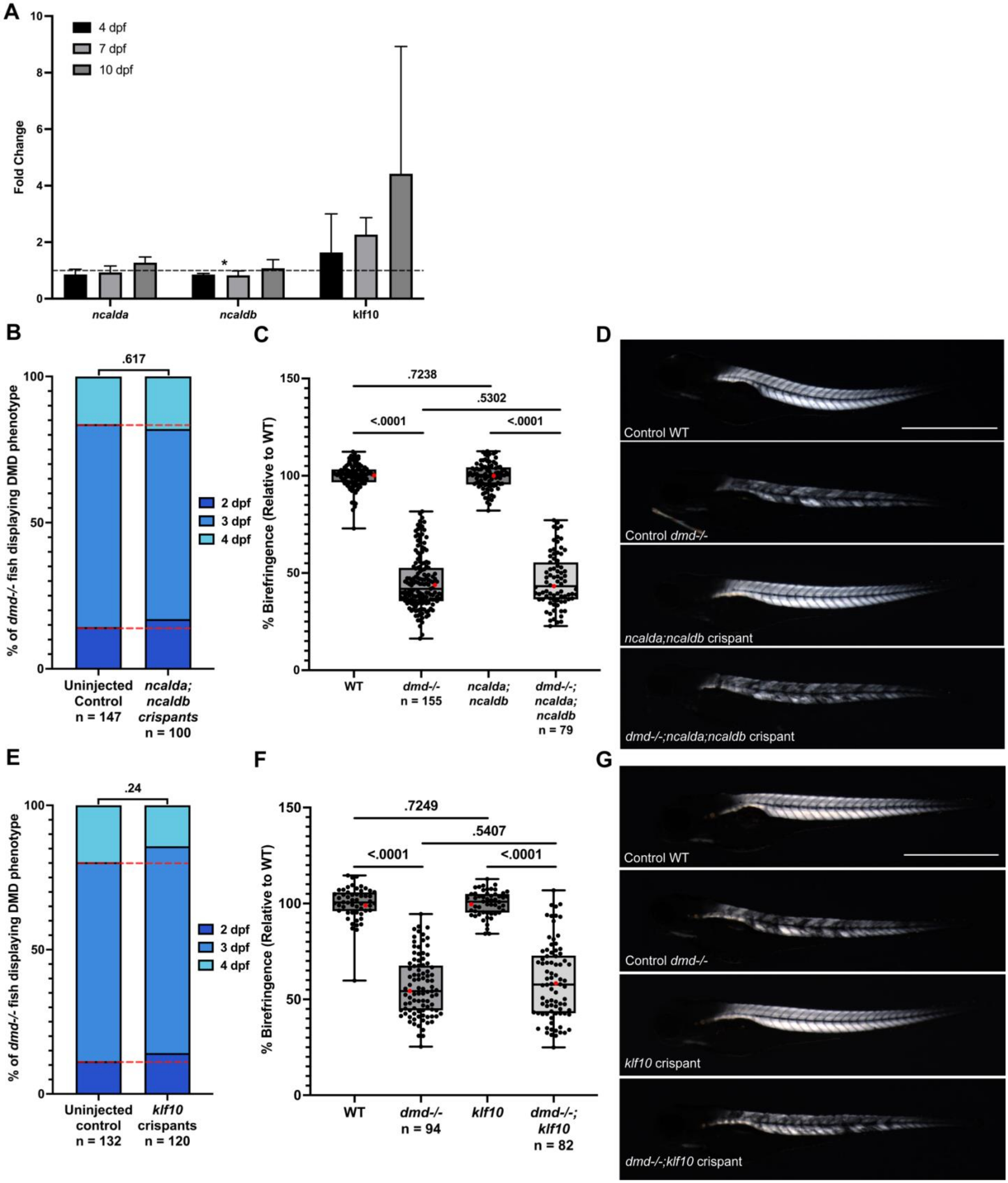


**Supplemental Figure 4: Mutation of *ncalda* and *ncaldb* or *klf10* does not have an impact on DMD onset or birefringence intensity.** (A) WT (Black) and *dmd-/-* mutants were raised to 4 (grey), 7 (dark grey), and 10 dpf (light grey) and processed for qPCR analysis, using 3 biological replicates (n=15 fish) for each sample and time point. qPCR data are presented as mRNA fold change in *dmd-/-* mutants relative to WT siblings of the same age, with the black dashed line indicating equivalent WT expression. *eef1a1a* was used as the normalization gene. Welch's T-Test used, * denotes significant difference. (B-G) Embryos from a *dmd+/-* intercross were injected with *ncalda* and *ncaldb* (B-D) or *klf10* (E-G) targeting CRISPR(s) or raised as uninjected controls. (B,E) DMD onset was assessed at 2 dpf (dark blue), 3 dpf (blue), and 4 dpf (light blue). Chi-square analysis was carried out to determine if there was a significant different in onset age proportions. (C,F) Box plots show overall birefringence intensities. Mann-Whitney U-Tests carried out for comparisons of overall birefringence intensities. Red dots denote representative images shown in panels D,G. (D,G) Representative birefringence images for each genotype. Scale bar denotes 1mm. Experiments were conducted at minimum in biological triplicate. Sample sizes listed below each genotype for the given experiment.

| Gene | F Primer | R Primer |
|---|---|---|
| *etaa1a* | AGCGGTGGATATATCTGACATTG | CCACTGCAGTAAAGAAGAGTCC |
| *etaa1b* | ATCCCACCATTGCACGTT | AGCTGAACACTTGGGTGTC |
| *ncalda* | GAAACGGCTACATCAGCAAATC | TCCTCTGGCATCTTCATCAC |
| *ncaldb* | GACGGCAACGGATACATCA | GACTCGTCCTCTGGCATTT |
| *man1a1* | GAGCAACAGGGCACGAG | CCAGGCGTAGTGCTTGTAAT |
| *galntl6* | GAGAAGAGACTGGCATGACTATG | ATCCTCAACAAGCGGGTATG |
| *adamts17* | CTAAACATCCGAGTCACCAAGC | CCAGTGACAGAAGCTCTCCA |
| *ltbp4* | GAGAACGGACAGGAGGAATTT | GCTGGGAAATACCGTAGTAACA |
| *pard6ga* | ACAGTCTTTACTGCGCATCTT | GGCTTATGGATTTCTTGCGTTT |
| *pard6gb* | CCCTGCTGCGGATCTTTAT | GAAGAGTGACCACTGCCTTT |
| *klf10* | AGCATGTCATCTCCATCTCACA | GTGGTTTGACCTGTGACCTCT |
| *thbs1a* | TGGAGACGGGACACACT | GTGTTCTCACACTGATGGACTC |
| *thbs1b* | CAATCCACACCATCCTGACT | AGCGTGGTTCCGAATACAA |
| *eef1a1a* | TTCTCCGAGTATCCTCCTCTG | CTTCTCCACTCCTTTAATCACTCC |

**Supplemental Table 1: Primers utilized for qPCR analysis.**

| Gene | CRISPR |
| --- | --- |
| ***thbs1a*** | TCATGGCTGCCGTCAGCACA |
| ***thbs1b*** | GCTGGTGTATGTCTTCACAA |
| ***lbp4*** | CGTACACTCGTTGATTCCCC |
| ***etaa1a*** | CATTGCTGGGGGAGTCATCG |
| ***etaa1b*** | ATCCCAGATGATATCTTGCG |
| ***man1a1*** | CGTGTCCAGTGCGTCCACAA |
| ***adamts17*** | CTCCAGTATAGAAACACCGG |
| ***ncalda*** | CCGACAAATGGACACCAATC |
| ***ncaldb*** | CCTTCGACGCCAATGGAGAT |
| ***galntl6*** | TGTGCGCACTAAGAAGCGAG |
| ***klf10*** | CGGATCACACTGATGGCCTG |

**Supplemental Table 2: CRISPR guide sequences.**

| Gene | HRMA F | HRMA R |
|---|---|---|
| ***thbs1a*** | TGGGGTTAAATTGTTTTCTCGGCAGG | TGCCCAGGTAGATGCAGTTGGC |
| ***thbs1b*** | TGCAGACGGATGACAAAAACATGC | GGGCTCATACCTGGCAAGTGCA |
| ***lbp4*** | GGGTCTGCGCAGTCAGGAGATG | AGGGTTTGTGCTCAGCCTGTGT |
| ***etaa1a*** | TCCAACACGTCCAGCAAGAGGT | AGCAGACACACGTACCATGCCG |
| ***etaa1b*** | CGGTTCCTACGGTGGAGATTCGC | TGCCATTCAAATCTTGCGTGGGT |
| ***man1a1*** | CCCCACAGGATGCCTTCTCAAA | CACCCATTCGGTAGCGGCTTCA |
| ***adamts17*** | GAGGAGGGCAGTGTTCAGC | TACAGCAGTTTCCACCACTCAC |
| ***ncalda*** | TGCCAGAGGATGAGTCCACACCT | TGAAAGTGGGAGAAAAAGGACCA |
| ***ncaldb*** | CGAGTTCAAGAAGATCTACGGC | ATGTTACACTCAGCGCAATGAT |
| ***galntl6*** | AGAGCACCTGAAGGCACATC | AAAAGTCAGGACCTCTCCTCTG |
| ***klf10*** | TCCACCGTCCAGCCCTGATCAG | GCTGACGGGAGATCTGGGACGT |

**Supplemental Table 3: Primers used for HRMA following CRISPR microinjection.**
